# COVID-19-associated olfactory dysfunction reveals SARS-CoV-2 neuroinvasion and persistence in the olfactory system

**DOI:** 10.1101/2020.11.18.388819

**Authors:** Guilherme Dias De Melo, Françoise Lazarini, Sylvain Levallois, Charlotte Hautefort, Vincent Michel, Florence Larrous, Benjamin Verillaud, Caroline Aparicio, Sebastien Wagner, Gilles Gheusi, Lauriane Kergoat, Etienne Kornobis, Thomas Cokelaer, Rémi Hervochon, Yoann Madec, Emmanuel Roze, Dominique Salmon, Hervé Bourhy, Marc Lecuit, Pierre-Marie Lledo

## Abstract

While recent investigations have revealed viral, inflammatory and vascular factors involved in SARS-CoV-2 lung pathogenesis, the pathophysiology of neurological disorders in COVID-19 remains poorly understood. Yet, olfactory and taste dysfunction are rather common in COVID-19, especially in pauci-symptomatic patients which constitutes the most frequent clinical manifestation of the infection. We conducted a virologic, molecular, and cellular study of the olfactory system from COVID-19 patients presenting acute loss of smell, and report evidence that the olfactory epithelium represents a highly significant infection site where multiple cell types, including olfactory sensory neurons, support cells and immune cells, are infected. Viral replication in the olfactory epithelium is associated with local inflammation. Furthermore, we show that SARS-CoV-2 induces acute anosmia and ageusia in golden Syrian hamsters, both lasting as long as the virus remains in the olfactory epithelium and the olfactory bulb. Finally, olfactory mucosa sampling in COVID-19 patients presenting with persistent loss of smell reveals the presence of virus transcripts and of SARS-CoV-2-infected cells, together with protracted inflammation. Viral persistence in the olfactory epithelium therefore provides a potential mechanism for prolonged or relapsing symptoms of COVID-19, such as loss of smell, which should be considered for optimal medical management and future therapeutic strategies.

## Introduction

COVID-19, caused by SARS-CoV-2 commonly induces airway and pulmonary symptoms, and in severe cases leads to respiratory distress and death (1). While COVID-19 is primarily regarded as a respiratory disease, many patients exhibit extra-respiratory symptoms of various severity. Among these, a sudden loss of olfactory function in SARS-CoV-2-infected individuals was reported worldwide at the onset of the pandemic. Loss of smell (anosmia) and/or of taste (ageusia) are considered now as cardinal symptoms of COVID-19 (2–4). Likewise, a wide range of central and peripheral neurological manifestations have been observed in severe patients. Although neuropilin-1 was recently found to facilitate SARS-CoV-2 entry in neural cells (5), and thus a neurotropism of SARS-CoV-2 could be suspected, a direct role of the virus in the neurological manifestations remains highly debated (2, 6).

The *bona fide* virus entry receptor is the angiotensin-converting enzyme 2 (ACE2), which is expressed along the entire human respiratory system, thereby accounting for SARS-CoV-2 respiratory tropism (7, 8). In the upper airways, in the superior-posterior portion of the nasal cavities resides the olfactory mucosa. This region is where the respiratory tract is in direct contact with the central nervous system (CNS), via olfactory sensory neurons (OSN), of which their cilia emerge within the nasal cavity and their axons project into the olfactory bulb (9). As loss of smell is a hallmark of COVID-19 and several respiratory viruses (influenza, endemic human CoVs, SARS-CoV-1) invade the CNS through the olfactory mucosa via a retrograde route (10), we hypothesized that SARS-CoV-2 might be neurotropic and capable of invading the CNS through OSNs.

SARS-CoV-2 can infect neurons in human brain organoids (11) and recent reports have confirmed the presence of SARS-CoV-2 in olfactory mucosa OSNs that express neuropilin-1 (5) and deeper within the CNS at autopsy (12, 13). Yet, the portal of entry of SARS-CoV-2 in the CNS remains elusive, as well as the exact mechanism leading to the olfactory dysfunction in COVID-19 patients. Various hypotheses have been proposed such as conductive loss due to obstruction of the olfactory cleft (14), alteration of OSN neurogenesis (15) and secondary CNS damage related to edema in the olfactory bulb (16, 17). Detailed study of the olfactory system and olfaction in living COVID-19 patients is thus needed to investigate the SARS-CoV-2 neuroinvasiveness in the olfactory epithelium.

Complementary to this approach, animal models recapitulating the biological and clinical characteristics of SARS-CoV-2-related anosmia would constitute useful tools to address deeper mechanisms. In this regard, wild-type mice are poorly susceptible to SARS-CoV-2 infection as the mouse ACE2 ortholog is not acting as a receptor for this virus (18), and the various transgenic mouse lines expressing the human version of the virus entry receptor (hACE2) under the control of different promoters, display disproportionate high-levels of CNS infection leading to fatal encephalitis (19–22), which rarely occurs in COVID-19 patients. This mismatch likely reflects the artefactual ectopic and high level of hACE2 expression caused by the different transgene promoters. In contrast, the golden Syrian hamster *(Mesocricetus auratus)* expresses an endogenous ACE2 protein able to interact with SARS-CoV-2 (18) and constitutes a naturally-permissive model of SARS-CoV-2 infection (23–25). Previous reports have shown infection in hamster olfactory mucosa, but whether olfactory neurons can be infected or only non-neuronal, epithelial sustentacular cells, is still controverted (26, 27). Moreover, the link between infection, neuroinflammation and tissue disruption of the olfactory neuroepithelium is unclear. Likewise, how damage of the neuroepithelium correlates with anosmia, and the potential SARS-CoV-2 neuroinvasion from the olfactory system to its downstream brain structures, remains highly debated.

Here, we report the interactions of SARS-CoV-2 with the olfactory system and its pathophysiological mechanisms. We first investigated SARS-CoV-2 infection of the olfactory mucosa in COVID-19 patients with recent loss of smell. Because olfactory mucosa biopsy is an invasive procedure, which cannot be used for research purpose in COVID-19 patients, we performed nasal mucosa brush sampling, a non-invasive technique previously used in patients to study neurodegenerative and infectious diseases (28–30). We next attempted to model SARS-CoV-2-associated anosmia/ageusia in golden Syrian hamsters to further investigate the pathogenesis of neuroepithelium and CNS infection. Finally, we investigated the olfactory mucosa of post-COVID-19 patients presenting long-lasting olfactory dysfunction.

## Results

### SARS-CoV-2 detection in the olfactory mucosa of COVID-19 patients with acute olfactory function loss

We enrolled 5 patients that were referred to the ear, nose and throat (ENT) department for olfactory function loss and COVID-19 suspicion in the context of the COVID-19 first wave in Paris, France, alongside with 2 healthy controls. The main clinical features of patients and controls are listed in Table 1. The time from first COVID-19 related symptoms to inclusion in the study ranged from 1 to 13 days. None of the patients required hospitalization. Their prominent symptom was recent loss of olfactory function (sudden for 4 patients but progressive for case #1) and was accompanied with taste changes (except case #3) and at least one symptom belonging to the clinical spectrum of COVID-19, such as diarrhea, cough, dyspnea, conjunctivitis, fever, fatigue, headache, muscle pain, laryngitis or a sore throat (Supplemental Figure S1A). Olfactory function loss was the first symptom related to COVID-19 in case #5 while it was preceded by, or concomitant with other symptoms in the remaining patients. Smell loss was deemed severe for cases #1, #2, #4 and #5, and moderate for case #3. Taste loss was deemed severe for cases #1, #2, #4 and #5, and mild for case #3. The characteristics of the taste and smell abnormalities are listed in Supplemental Table S1. Other otolaryngologic symptoms were rhinorrhea for 4 patients, not concomitant with smell loss, nasal irritation for 2 patients and hyperacusis for case #1. Nasal obstruction was not reported in any of the patients. Taste changes were characterized in the 4 patients by dysgeusia where they had a reduced acuity for sweet taste, had a bad taste in the mouth, reduced or increased acuity for bitter, reduced acuity for salt or sour were reported in 3 out of the 4 patients with dysgeusia. Two patients (#2 and #4) were unable to discriminate between different foods such as meat and fish.

**Table 1.** Features at inclusion of the participants with recent loss of smell associated to COVID-19.

| Patient/Control | COVID #1 | COVID #2 | COVID #3 | COVID #4 | COVID #5 | Control #1 | Control #2 |
| --- | --- | --- | --- | --- | --- | --- | --- |
| **Years/Sex** | 53/W | 31/W | 61/W | 40/W | 46/M | 31/W | 47/M |
| **Neurological features at inclusion** | Anosmia-Ageusia-Hyperacusis | Anosmia-Phantosmia-Ageusia | Hyposmia | Anosmia-Parosmia-Ageusia | Anosmia-Ageusia | - | - |
| **Comorbidities<sup>a</sup>** | Overweight | - | Diabetes-Hypertension | - | - | - | - |
| **Smoking status** | Nonsmoker | Previous<br>(5 years ago) | Nonsmoker | Current | Nonsmoker | Nonsmoker | Nonsmoker |
| **Time between the first disease symptoms<sup>b</sup> and inclusion** | 7 | 8 | 2 | 1 | 13 | - | - |
| **Time between the first disease symptoms<sup>b</sup> and olfactory loss** | 2 | 5 | 0 | 0 | 0 | - | - |
| **SARS-CoV-2 PCR in the nasopharynx** | Neg | Pos | Neg | Neg | Pos | Not done | Neg |
| **SARS-CoV-2 PCR in the olfactory mucosa<sup>c</sup> (copy number/μL<sup>d</sup>)** | Pos (<200) | Pos (2.25. 10<sup>6</sup>) | Pos (<200) | Pos (<200) | Pos (<200) | Neg | Neg |
| **SARS-CoV-2 antigens in the olfactory mucosa** | Yes | Yes | Yes | No | No | No | No |
| **IBA-1 positive cells in the olfactory mucosa** | Yes | Yes | Yes | Yes | Yes | No | No |
| **IL-6 RNA in the olfactory mucosa (log<sub>2</sub> (2ddct))** | 5,41 | 7,69 | 4,33 | -0,67 | 1,3 | 0 | 2,79 |
| **CXCL10 RNA in the olfactory mucosa (log<sub>2</sub> (2ddct))** | 3.38 | 6.97 | 1.51 | 1.65 | 0.85 | 0 | -4.35 |
| **CCL5 RNA in the olfactory mucosa (log<sub>2</sub> (2ddct))** | -0.80 | -0.27 | -0.80 | 0.17 | 1.02 | 0 | 3.96 |
| **ISG20 RNA in the olfactory mucosa (log<sub>2</sub> (2ddct))** | -1.88 | -0.98 | 0.41 | 0.66 | 1.06 | 0 | 2.88 |
| **Mx1 RNA in the olfactory mucosa (log<sub>2</sub> (2ddct))** | -4.31 | 0.53 | 2.18 | 2.71 | 3.46 | 0 | 0.45 |
| **Smell changes scores<sup>e</sup>**<br>**Score range: 0-5** | 5 | 5 | 3 | 4 | 4 | 0 | 0 |
| **Taste changes scores<sup>e</sup>**<br>**Score range: 0-10** | 6 | 10 | 1 | 10 | 9 | 0 | 0 |
| **Combined taste & smell changes scores<sup>e</sup>**<br>**Score range: 0-15** | 11 | 15 | 4 | 14 | 13 | 0 | 0 |
| **Visual analogue scale score for smell<sup>f</sup>**<br>**Score range: 0-100** | 5 | 2 | 45 | 11 | 95 | 100 | 100 |
| **Visual analogue scale score for taste<sup>g</sup>**<br>**Score range: 0-100** | 6 | 5 | 100 | 0 | 75 | 100 | 100 |

W: woman; M: man; <sup>a</sup>Overweight: 25-29.9 kg/m<sup>2</sup>; <sup>b</sup>Two or more of the following signs or symptoms: Fever, cough, headache, fatigue, muscle pain, sore throat, diarrhea, conjunctivitis, laryngitis; dyspnea, anosmia; <sup>c</sup>RT-PCR Sybr; <sup>d</sup>RT-PCR Taqman; <sup>e</sup>TTS: Taste and Smell Survey: Sum of smell changes scores and taste changes scores; Smell changes scores from 0 normal to 5 anosmia; Taste changes scores from 0 normal to 10 ageusia; <sup>f</sup>VAS: Visual Analogue Scale from 0 (anosmia or ageusia) to 100 normal. Five shades of gray are used for scoring from 0 no color to maximum dark.

To investigate whether infection in the olfactory mucosa is associated with olfactory functional loss, all patients underwent olfactory mucosa brush cytological sampling. Only two patients had detectable SARS-CoV-2 RNA, using the conventional nasopharyngeal samples at inclusion (Table 1). However, all patients – but none of the controls – had detectable SARS-CoV-2 RNA in cytological samples from the olfactory mucosa using the RT-qPCR SYBR green technique, unambiguously confirming the diagnosis of COVID-19 (Table 1). Patient #2 had a strong viral load in the olfactory mucosa (2.25. 10^6^ RNA copies/μL), while other cases were positive (RT-qPCR SYBR green) but not quantifiable (less 200 RNA copies/μL using the RT-qPCR Taqman technique).

We further investigated the viral presence in the patient’s olfactory mucosa by immunofluorescence labeling of the cytological samples. Variable cell density between olfactory mucosa samples from the COVID-19 and control patients was found, but all samples contained mature OSNs, positive for OMP, validating the quality of the swabbing procedure (Figure 1, A and B and Supplemental Figure S1B). Immunostaining revealed the presence of SARS-CoV-2 antigens (nucleoprotein, NP) in 3 patients (RT-qPCR+) out of 5 but not in controls (Table 1, Figure 1). We observed numerous Iba1^+^ cells in the olfactory mucosa of all patients whereas few to no Iba1^+^ cells in controls (Table 1; Figure 1E and Supplemental Figure S1D). These data suggest that SARS-CoV-2 infection is associated with inflammation of the olfactory mucosa in patients with olfactory impairment, thus we measured the profile of local cytokine and inflammatory mediators (Table 1). Whereas there was no change in the transcript levels of *Ccl5, Cxcl10, Isg20* and *Mx1* genes as compared to controls, transcript levels for the proinflammatory cytokine *IL6* were elevated in the 3 patients with detectable SARS-CoV-2 antigens as compared to control patients, and the 2 other patients positive for CoV-2 RNA but without detectable CoV-2 antigens (Table 1).

**Figure 1–.**
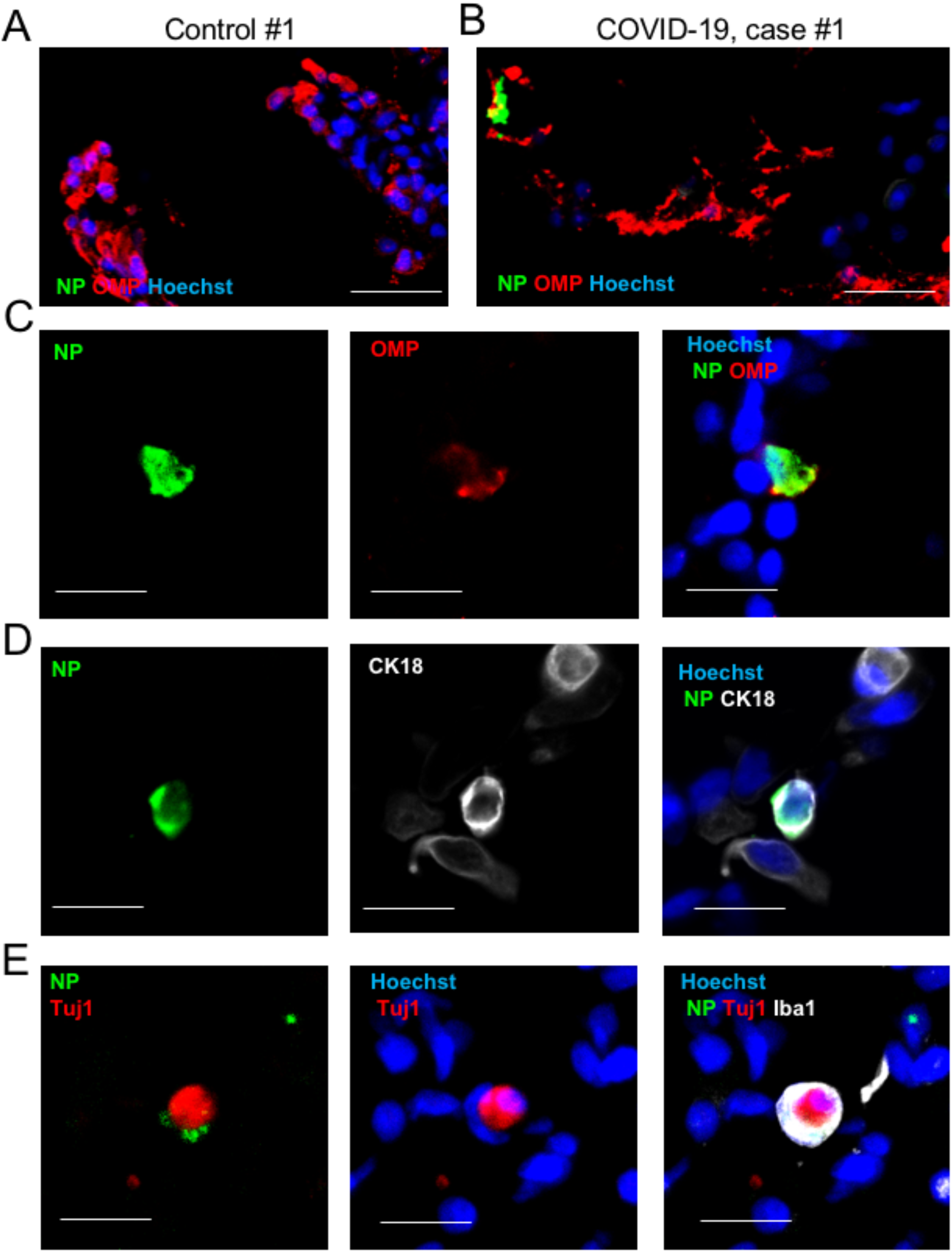
Analysis of olfactory mucosa from COVID-19 patients with acute olfactory function loss, at early stage of infection. (**A**) Immunofluorescence of cells retrieved from the olfactory mucosa of the control subject #1. (**B**) Cells retrieved from the olfactory mucosa of the COVID-19 patient #1. (**C-E**) Close-up immunofluorescence images of olfactory epithelium samples from the COVID-19 patient #2. Infected mature olfactory neurons (OMP^+^) are observed (C), alongside sustentacular CK18^+^ cells (D) and Iba1^+^ myeloid cells engulfing Tuj1^+^ neuron parts (E). SARS-CoV-2 is detected by antibodies raised against the viral nucleoprotein (NP). Scale bars = 20μm (A, B) or 10μm (C-E).

Together, this first set of data indicates that SARS-CoV-2 exhibits a clear tropism for the olfactory epithelium, and this infection is associated with increased local inflammation. We next investigated the identity of the cell types targeted by SARS-CoV-2. We detected SARS-CoV-2-infected mature sensory neurons (OMP^+^; Figure 1, B and C); other SARS-CoV-2 infected cells were sustentacular cells (expressing CK18, see Figure 1D and Supplemental Figure S1C), and myeloid cells (expressing Iba1, Figure 1E). We detected the presence of several immature sensory neurons (Tuj-1^+^) in the olfactory mucosa of all patients, some of them being infected. Interestingly, some Iba1 and SARS-CoV-2 positive cells were engulfing portions of Tuj-1 cells in the olfactory mucosa of COVID case #2, suggesting that infected immature sensory neurons were in the process of being phagocytosed by brain innate immune cells (Figure 1E). These results show that a variety of cell types are infected in the olfactory epithelium of COVID-19 patients. Among them, the mature OSN are critically relevant in the context of the anosmia. To assess the impact of the neuroepithelium infection by SARS-CoV-2, we infected Syrian golden hamsters to experimentally reproduce anosmia and ageusia, and investigate the potential SARS-CoV-2 infection of the olfactory system and upstream brain tissues.

### Modeling SARS-CoV-2 taste and smell function loss using nasal instillation in golden hamsters

Syrian golden hamsters (both sexes) were intranasally inoculated with 6.10^4^ PFU of SARS-CoV-2 and followed-up for several days. Clinical, sensorial and behavioral functions were assessed at different timepoints (Supplemental Figure S2A). SARS-CoV-2 inoculation resulted in a decrease in body weight and a degradation in the clinical score as early as 2 days post-inoculation (dpi), with a peak between 4 and 6 dpi, and sickness resolution by 14 dpi (Figure 2, A and B). High viral loads were detected throughout the airways of infected hamsters at 2 and 4 dpi and remained detectable even at 14 dpi (Figure 2C), consistent with the well-established respiratory tropism of SARS-CoV-2. In line with our observations in human samples, the nasal turbinates of infected hamsters exhibited high viral loads as soon as 2 dpi. Strikingly, viral RNA was also detected from 2 dpi in various parts of the brain, including the olfactory bulb, cerebral cortex, brainstem (diencephalon, midbrain, pons and medulla oblongata) and cerebellum. The olfactory bulb was by far the most infected brain region, from 2 dpi and onward (Figure 2D). Having shown the concomitant infection of nasal turbinates and the CNS, we further investigated their impact on sensory and behavioral responses.

**Figure 2–.**
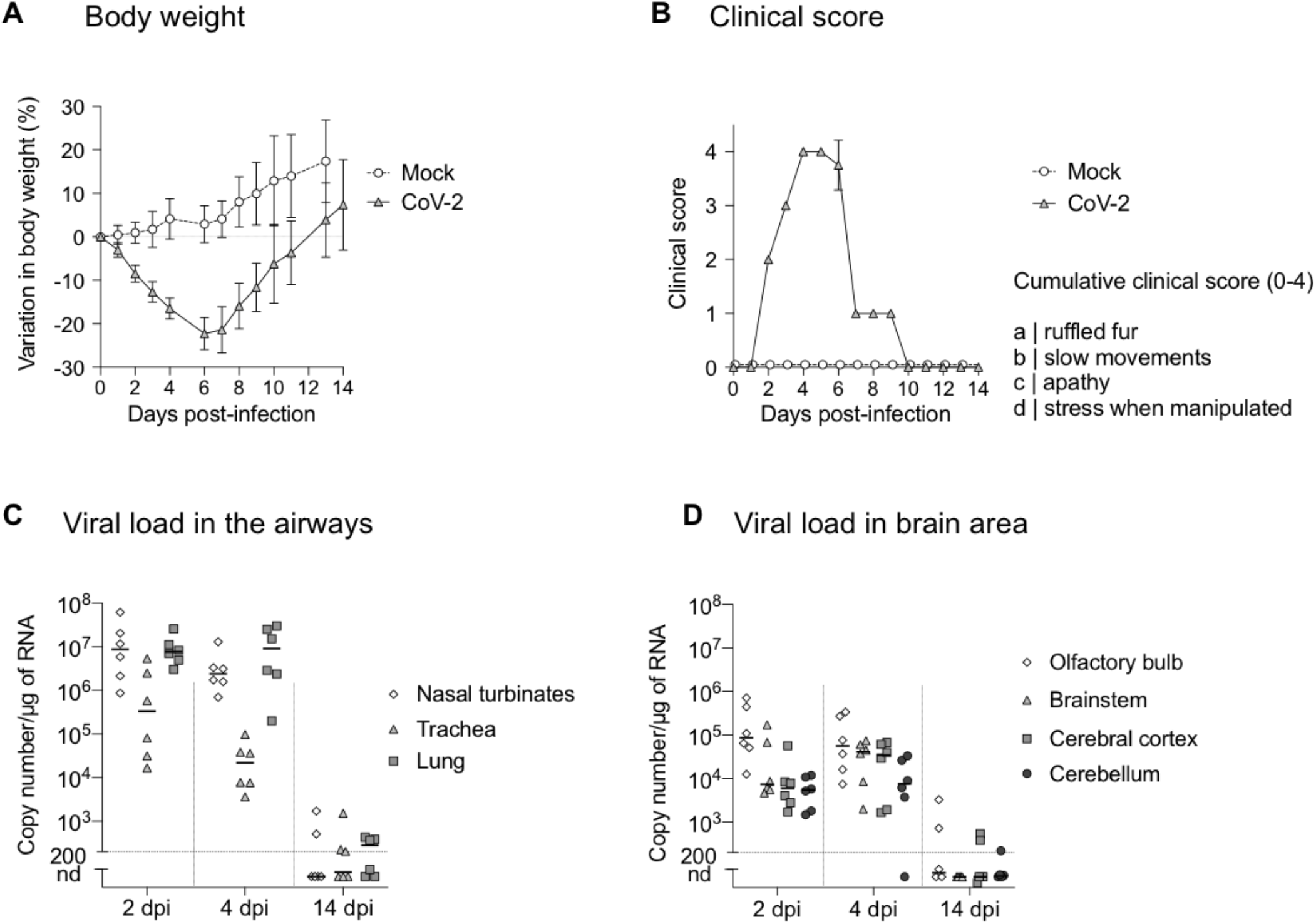
Clinical and molecular characteristics of experimental infection with SARS-CoV-2 in golden hamsters. (**A-B**) Variation in body weight (**A**) and clinical score (**B**) of mock-and SARS-CoV-2 infected hamsters for 14 days post-infection (dpi). (**C, D**) Quantification of SARS-CoV-2 RNA in hamster airways (**C**) and in different brain areas (**D**) at 2, 4 and 14 dpi. Horizontal lines indicate medians. N=4-8/ timepoint in (**A, B**); N=6/timepoint in (**C, D**).

We assessed both gustatory and olfactory function of SARS-CoV-2-inoculated hamsters. At 2 dpi, we subjected hamsters to a sucrose preference test. As expected, mock-infected animals displayed a clear preference towards sucrose-complemented water *vs.* control water, whereas infected hamsters had no preference towards the sucrose-complemented water (Figure 3A), indicative of a SARS-CoV-2-associated dysgeusia/ageusia (and/or sickness-induced anorexia/anhedonia). Moreover, infected animals exhibited signs of hyposmia/anosmia during food findings experiments, as they needed more time to find hidden (buried) food than uninfected hamsters, and a significant proportion of them (50% at 3dpi and 37.5% at 5 dpi) failed to find the food at the end of the test (Figure 3, B and C). Nevertheless, all infected hamsters succeeded to find visible food (Figure 3C) demonstrating that no sickness behavior, visual impairment or locomotor deficit accounted for the delay in finding the hidden food.

**Figure 3–.**
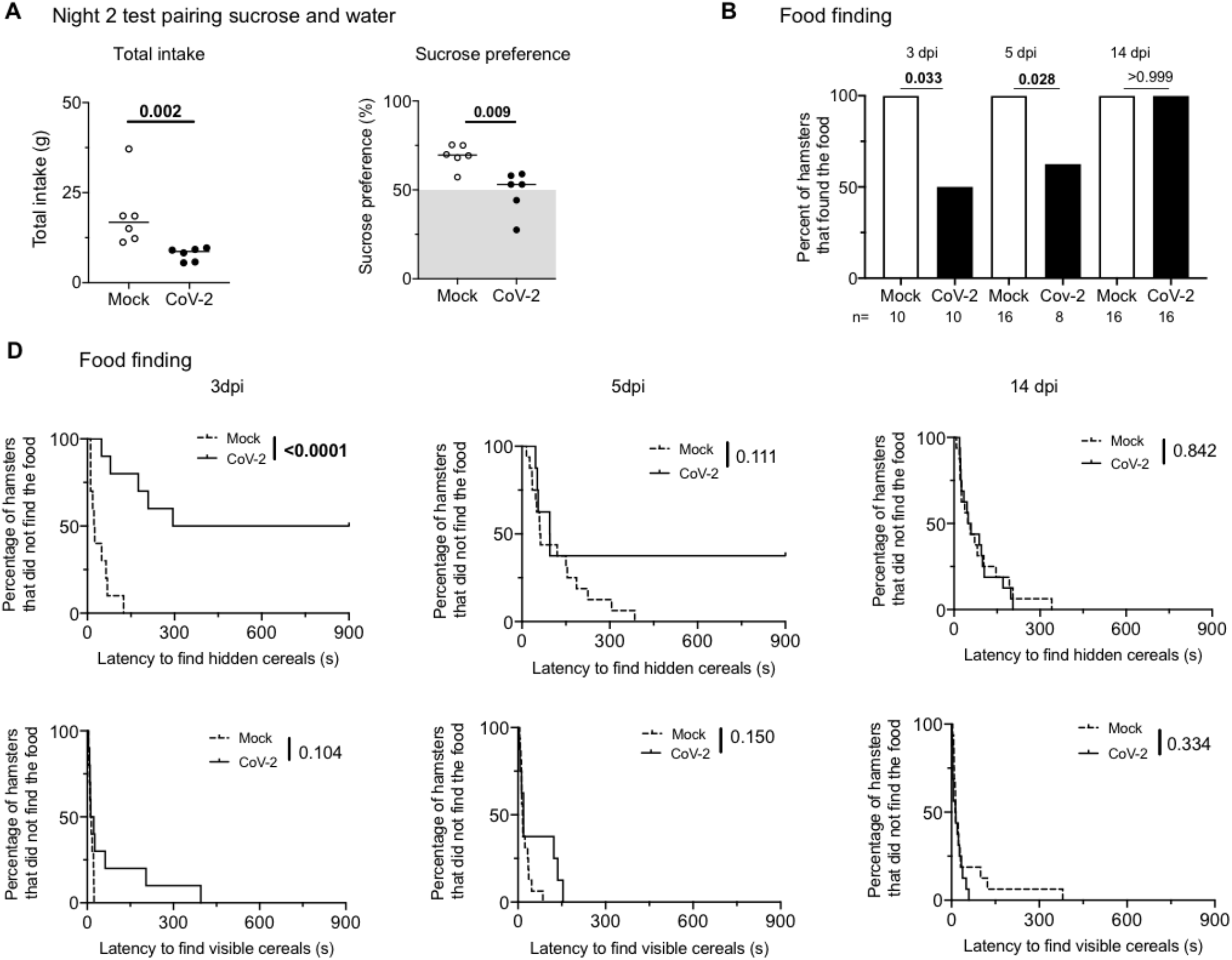
Experimental infection with SARS-CoV-2 in golden hamster induces transient anosmia and ageusia. (**A**) Variation in total consumption of liquid overnight and preference towards 2% sucrose-containing water of control and SARS-CoV-2 infected hamsters at 2 dpi. (**B**) Fraction of control or infected hamsters successfully finding hidden food in 15 minutes. (**C**) Fraction of control or infected hamsters successfully finding hidden or visible food over time. Food-finding assays were performed at 3, 5-and 14-dpi. Mann-Whitney test (A), Fisher’s exact test (B) and Log-rank (Matel-Cox) test (C). P value is indicated in bold when significant. Bars indicate medians. N=6-16 per group.

Also, no locomotor deficit was observed either during the open field (Supplemental Figure S2B) or painted footprint tests (Supplemental Figure S2C), further excluding a motor deficit bias during the food finding test. At 14 dpi, when weight and clinical score had resumed to standard levels (Fig. 2, A and B), all animals successfully found the hidden food, indicating that infection-associated anosmia recovered spontaneously in this animal model.

### SARS-CoV-2 promotes cellular damage in both upper and lower airways in infected hamsters

We then investigated the impact of SARS-CoV-2 infection on hamster olfactory mucosa which exhibited high viral loads (Figure 2C). The uppermost part of nasal turbinates is overlaid by the olfactory epithelium (Figure 4A), a neuroepithelium composed of sensory neurons and support sustentacular cells with both cell populations being ciliated. Imaging by scanning electron microscopy of the olfactory neuroepithelium showed an important loss of ciliation as early as 2 dpi (Figure 4, B and C) on large portions of the epithelial surface, indicating cilia loss in both OSNs and sustentacular cells (Supplemental Figure S3). At 4 dpi, viral particles were seen budding from deciliated cells (Figure 4D). At 14 dpi, the olfactory mucosa appeared ciliated anew, indistinguishable from that of mock-infected animals (Figure 4E and Supplemental Figure S3), consistent with the recovery of olfaction seen in infected hamsters (Figure 3C).

**Figure 4–.**
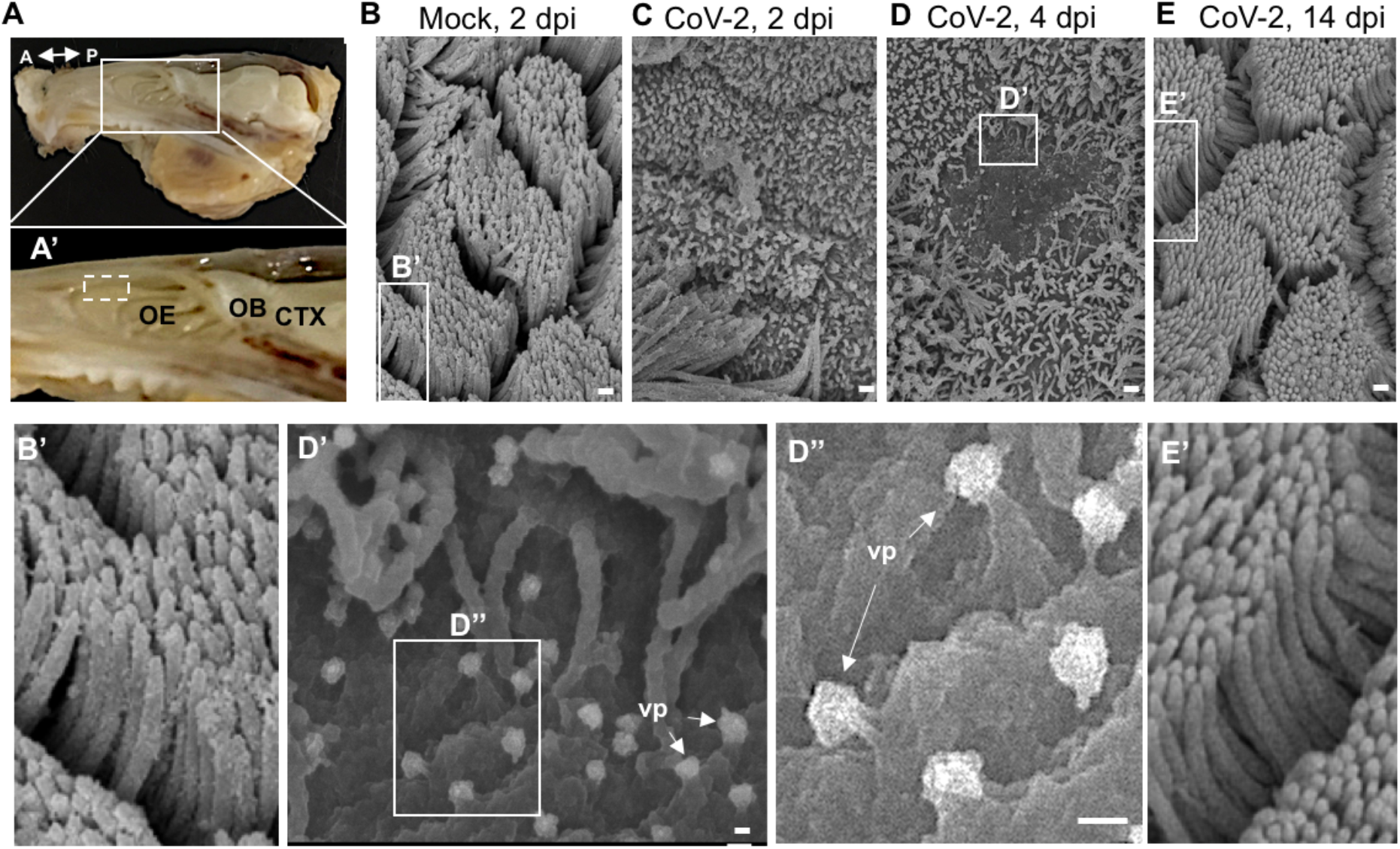
SARS-CoV-2 induces loss of ciliation in the olfactory epithelium. (**A**) Dissected hamster head, skin and lower jaw removed, sagitally cut in half. Double-headed arrow denotes the anteroposterior (A-P) axis. Close-up in **A’** shows the tight relationship between the olfactory epithelium (OE), the olfactory bulb (OB) and the cerebral cortex (CTX). Discontinuous square indicates the area collected for scanning electron microscopy. (**B-E**) Scanning electron microscope imaging showing changes in olfactory epithelium following CoV-2 infection. The olfactory epithelium of mock-(**B, B’**) and CoV-2 inoculated hamsters at 2 dpi (**C**), 4 dpi (**D, D’, D”**) and 14 dpi (**E, E’**). A loss of cilia is observed at 2 and 4 dpi for infected hamsters. Viral particles (vp) are seen emerging from deciliated cells (**D’-D’’**, white arrows). Scale bars: 1 μm (B-E), 100 nm (D’, D”).

In line with the detection of viral particles by electron microscopy at 4 dpi, SARS-CoV-2 immunostaining was detected in the hamsters’ olfactory mucosa at this time point which was associated with an infiltration of myeloid Iba1^+^ cells (Figure 5, B-E). In the olfactory mucosa, SARS-CoV-2 antigens were found in the cytoplasm of mature (OMP^+^; Figure 5B) and immature (Tuj 1^+^; Figure 5D) sensory neurons and in sustentacular cells (CK18^+^; Supplemental Figure S4B). Some Iba1^+^ immune cells seen infiltrating the neuroepithelium were positive for SARS-CoV-2, consistent with a potential secondary infection resulting from the phagocytosis of infected cells (Figure 5D, arrow). Of note, the areas of neuroepithelium containing infected cells were disorganized (see Figure 5, B and D, and Supplemental Figure S4B), while adjacent areas without SARS-CoV-2 remained stratified (Supplemental Figure S4C). Cilia of OMP^+^ neurons located at the apical part of olfactory epithelium were lost in the disorganized infected neuroepithelium (Supplemental Figure S4C).

**Figure 5–.**
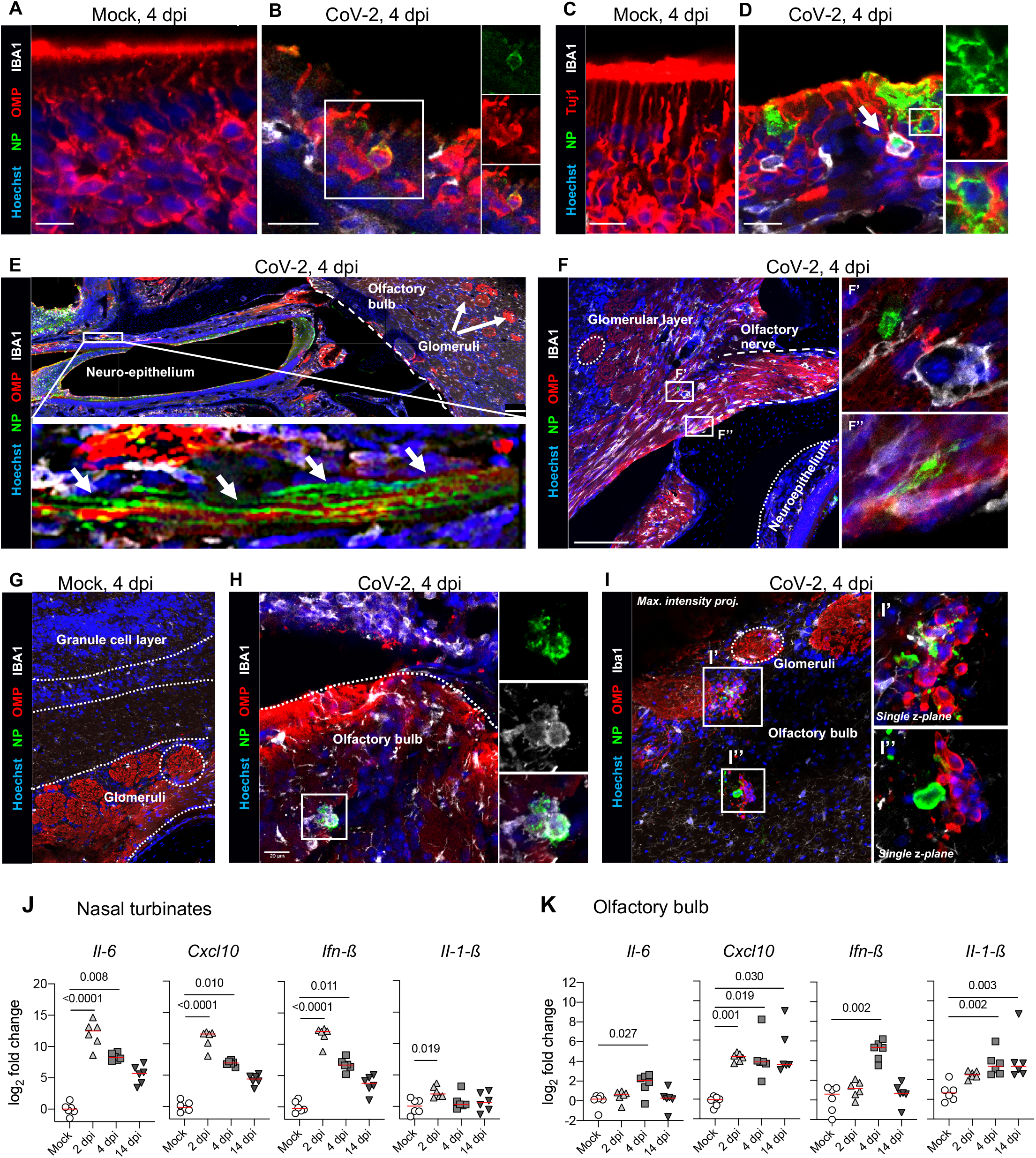
SARS-CoV-2 antigens detection and cytokine/chemokine transcripts quantification in the olfactory system of hamsters. (**A-D**) Olfactory epithelium of mock-(**A, C**) and SARS-CoV-2 (**B, D**) infected hamsters at 4 dpi. Insets show infected OMP^+^ mature olfactory sensory neurons (**B**), or infected Tuj1^+^ immature olfactory sensory neurons (**D**). The arrow in D indicates an infected Iba1+ cell. **E**) Sagittal section showing nasal turbinates and olfactory bulb of SARS-CoV-2 infected hamster at 4 dpi. Inset depicts the high density of SARS-CoV-2 staining in olfactory sensory neuron axons. (**F**) Olfactory sensory axons projecting into glomeruli in the olfactory bulb of SARS-CoV-2 inoculated hamsters at 4 dpi. Insets (**F’, F”**) show infected cells. (**G-I**) Olfactory bulb of mock-(**G**) or SARS-CoV-2 (**H, I**) infected hamsters at 4 dpi. Iba1^+^ infected cells are shown in (**H**) and several infected cells are observed in (**I**). SARS-CoV-2 is detected by antibodies raised against the viral nucleoprotein (NP). Scale bars: 20μm (A-D, G-I), 100μm (E, F). Images are single z-planes (A-H) or maximum intensity projection over a 6μm depth (I). (**J, K**) Cytokines and chemokines transcripts in the nasal turbinates (**J**) and in the olfactory bulb (**K**) at 2, 4 and 14 dpi. Kruskal-Wallis followed by the Dunn’s multiple comparison test (J, K). The p value is indicated when significant. Horizontal lines indicate the medians.

### SARS-CoV-2 dissemination to the brain and neuroinflammation in infected hamsters

Having shown that SARS-CoV-2 infects OSNs, and that SARS-CoV-2-infected hamsters exhibit signs of anosmia and ageusia, we wondered whether SARS-CoV-2 invades the CNS *via* a retrograde route from the olfactory system. SARS-CoV-2 was detected in olfactory nerve bundles close to the neuroepithelium, as demonstrated by the co-localization of SARS-CoV-2 nucleoprotein antigen and OMP^+^ sensory neuron axons reaching the olfactory bulb (Figure 5E), consistent with a retrograde infection of axons. Furthermore, SARS-CoV-2 nucleoprotein was detected at the junction of the olfactory nerve and olfactory bulb, seemingly infecting cells of neuronal/glial morphology (Figure 5F). In the olfactory bulb, SARS-CoV-2 nucleoprotein was detected in Iba1+ cells (Figure 5H) and in uncharacterized cells (Figure 5I) in the glomerular layer of the olfactory bulb. The viral nucleoprotein was not detected in other areas of the brain. The high viral RNA loads in the nasal turbinates and in the olfactory bulb, together with the observation of viral antigens along the entire route from the olfactory sensory organ to the bulb, suggests that SARS-CoV-2 enters the CNS through the olfactory system.

In the nasal turbinates, we detected an intense pro-inflammatory environment, with an upregulation of *Il6, Cxcl10, Ifnb1* and *Il1b* at 2 dpi, and a slight decrease at 4 and 14 dpi (Figure 5J). Similarly, the olfactory bulb exhibited an important upregulation in the expression of these genes (Figure 5K), but in a different and delayed pattern compared to the nasal turbinates: While *Cxcl10* was significantly overexpressed throughout the infection, the increase in *Il6*, *Ifnb1* and *Il1b* RNA levels was observed only at 4 dpi, with *Il1b* being up-regulated up to 14 dpi. These data reveal bulbar inflammation during SARS-CoV-2 infection, possibly in response to signaling via olfactory nerves.

Using RNA-seq, we observed 374 and 51 differentially expressed genes (DEG; increased or decreased, respectively) in the bulbs of SARS-CoV-2 infected hamsters at 4 dpi (Figure 6A). The DEG were classified according to KEGG (Kyoto Encyclopedia of Genes and Genomes) pathways (Figure 6B) and the GO (gene ontology) terms based on their biological processes, molecular functions and cellular components (Figure 6C). Upregulated genes were mainly involved in inflammatory responses and responses to virus infection, with innate immunity components (type-I IFN-mediated response, NK cell activation, TLRs, RLRs, NF-κB and Jak-STAT signaling pathways), adaptive immunity components (TH1, TH2, CD4^+^ T-cells) and functions related to chemokine signaling. Other biological processes related to nervous system functions were synapse pruning, upregulation of the neuroinflammatory response, and astrocyte and microglial activation. To validate the involvement of these signaling pathways, we analyzed the expression of selected targets in the olfactory bulb by RT-qPCR (Figure 6D). The genes *Mx2, Irf7, Ddx58* and *Stal1* gene transcripts were found significantly upregulated early in the infection (2 and 4 dpi), whereas *Ccl5* was upregulated only at 4 dpi. The overexpression of *Ccl5* and *Irf7* persisted even at 14 dpi. Altogether, SARS-CoV-2-associated inflammation in the bulb confirmed by unbiased RNA-seq analysis, along with the increased viral load detected in the brain parenchyma, supports the assumption that SARS-CoV-2 neuroinvasion drives neuroinflammation. Of note, *Cxcl10, Il1b, Ccl5* and *Irf7* overexpression persisted up to 14 dpi, when animals had recovered from ageusia/anosmia. These data indicate that an infectious or post-infectious inflammatory process persist even in the asymptomatic, or in a delayed post-symptomatic phase, in this animal model.

**Figure 6–.**
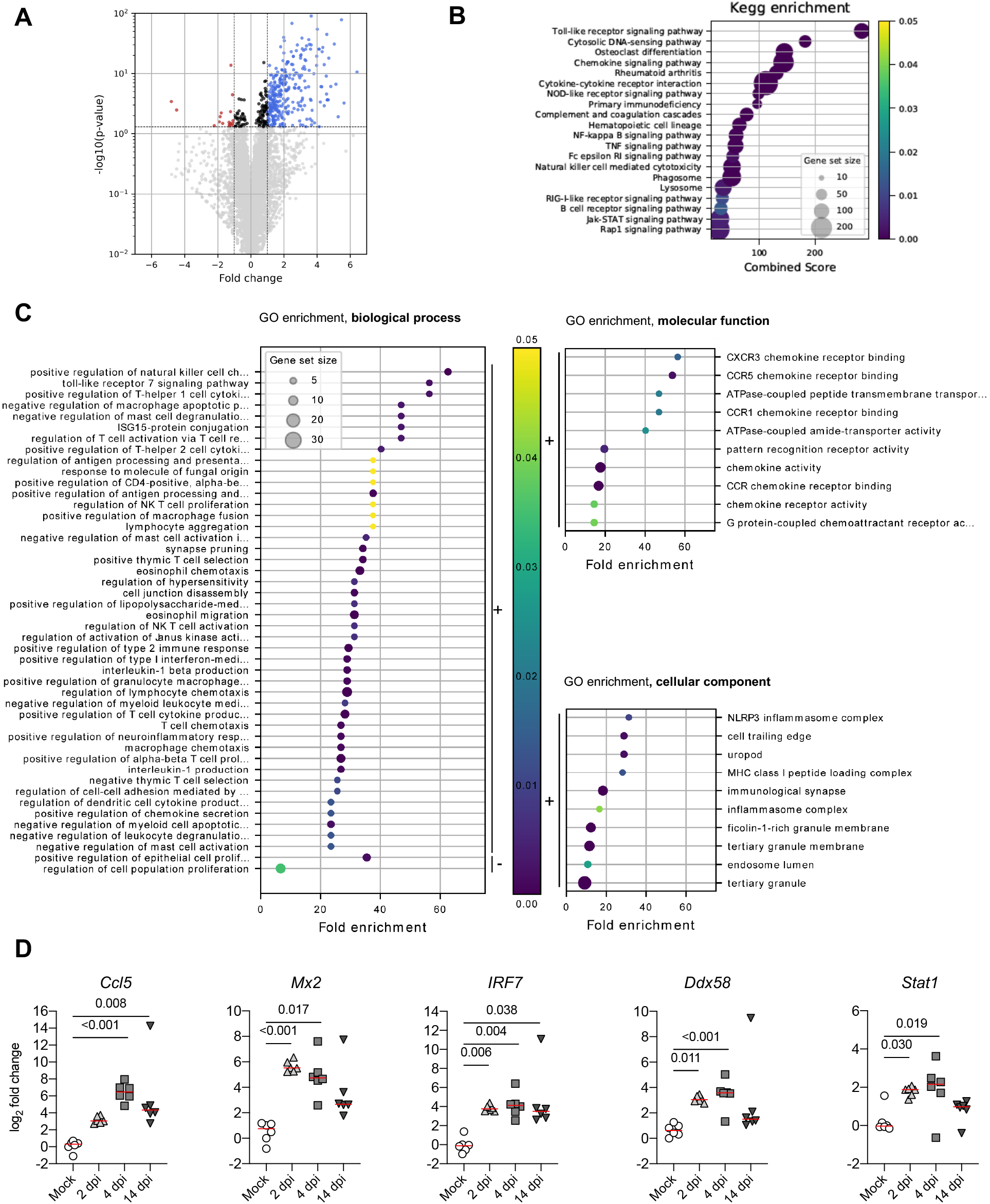
Differentially expressed genes in the olfactory bulb of golden hamsters infected by SARS-CoV-2 (at 4 dpi) derived by RNA-seq. (**A**). Volcano plot of the comparisons between infected and non-infected samples. Y-axis represents the Benjamini-Hochberg corrected p-value on a logarithmic scale (-log10). Grey dots represent genes not passing a threshold of FDR < 0.05. Black dots represent genes passing the FDR threshold but having fold changes between −1 and 1. Red and blue dots correspond to significant down and up-regulated genes with a fold change inferior to −1 or superior to 1, respectively. (**B**). KEGG-pathways enrichment based on the differentially regulated genes between infected and noninfected samples. Only the 20 highest combined scores are plotted. Circle sizes are proportional to the gene set size. Circle color is proportional to the corrected p-values and corresponds to the scale presented in C, D and E. (**C**). GO enrichment analysis considering biological process only. Selected GO terms are based on the up and down-regulated genes between infected and non-infected samples. The black bars on the right-hand side of the scatter plot indicate enrichment based on down (“-”) and up (“+”) regulated gene sets. Only the 50 highest fold enrichments are plotted for the up regulated gene set. Circle sizes are proportional to the gene set size, which shows the total size of the gene set associated with GO terms. Circle color is proportional to the corrected p-values and corresponds to the scale presented between C, D and E. GO enrichment analysis considering molecular function and cellular components related The figures follow the same construction as in biological process, with the exception that only the 10 highest fold enrichments are plotted for the up regulated gene set. (**D**) Validation targets in the olfactory bulb at 2, 4 and 14 dpi. n=6/time-point. Kruskal-Wallis followed by the Dunn’s multiple comparison test (J, K). The p value is indicated when significant. Horizontal lines indicate the medians.

### SARS-CoV-2 persistence in human olfactory mucosa with long-lasting/relapsing loss of smell

In some patients, neurological impairments and/or sensory dysfunctions persist even 3 months later from the onset of COVID-19 symptoms, and this may be linked to persistent viral infection and/or inflammation (31, 32). We recruited 4 patients seen with prolonged/recurrent olfactory function loss after COVID-19. The main characteristics of these patients are listed in Table 2. They were recruited in the cohort between July 15 and 29, 2020, at a time where viral circulation in Paris was very low (<10 cases/100,000 inhabitants/week), implying that SARS-CoV-2 reinfection of these patients was very unlikely. In this case, the time from first COVID-19 related symptoms to inclusion ranged from 110 to 196 days.

**Table 2.**
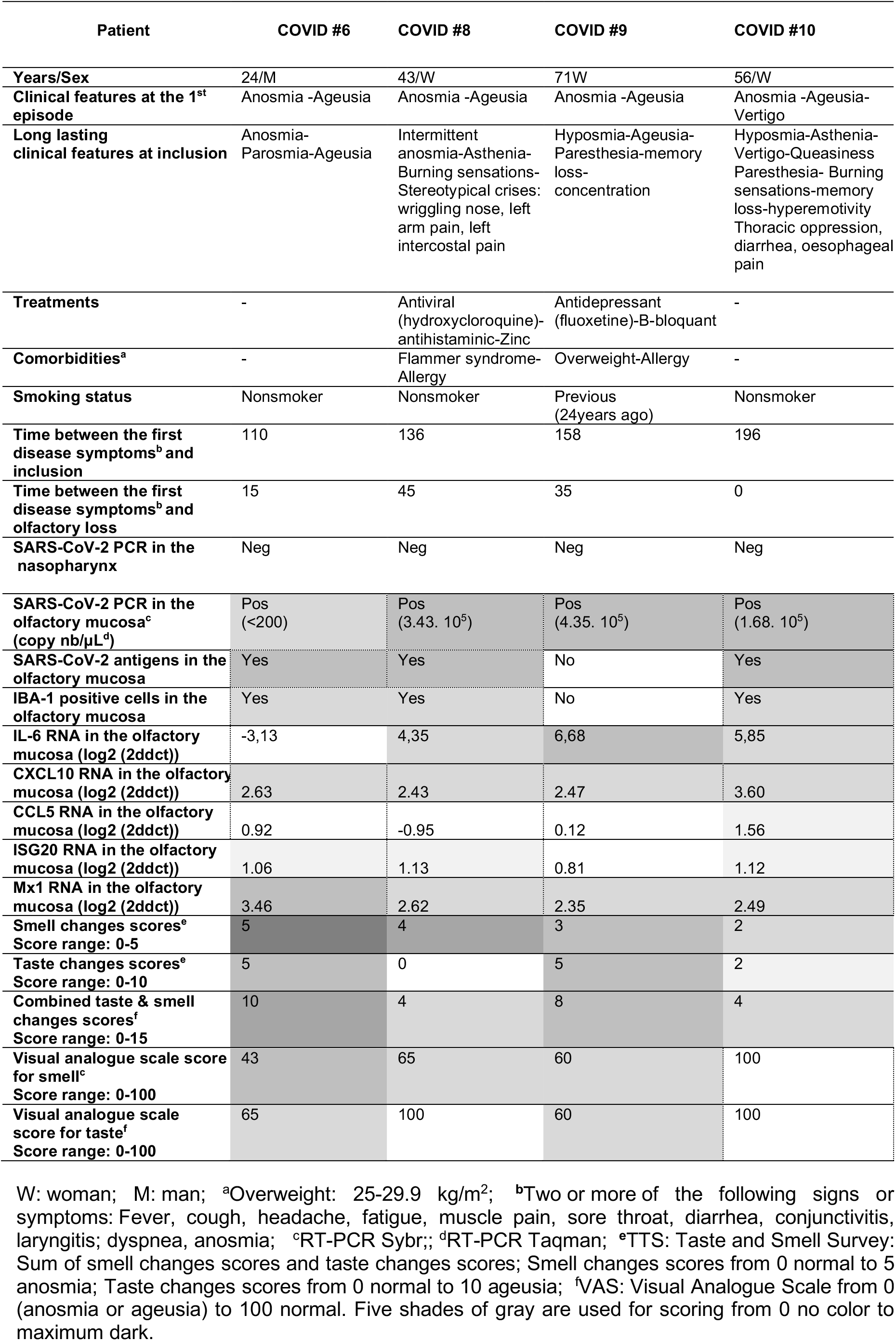
Individual features at inclusion of the participants with persistent olfactory dysfunction.

All patients have been previously diagnosed with COVID-19 between January and March 2020, based on their initial clinical assessment, including sudden anosmia at disease onset, accompanied with taste changes (except case #8) and at least one clinical sign related to COVID-19, such as fever, fatigue, diarrhea, cough, dyspnea, headache, muscle pain, laryngitis, sore throat, but also paresthesia and vertigo in some patients (Figure 7A). Smell loss was complete at disease onset for these patients. Other otolaryngologic symptoms were rhinorrhea for 2 patients, not concomitant with smell loss and nasal irritation for 3 patients. Nasal obstruction was reported in patient #10.

**Figure 7–.**
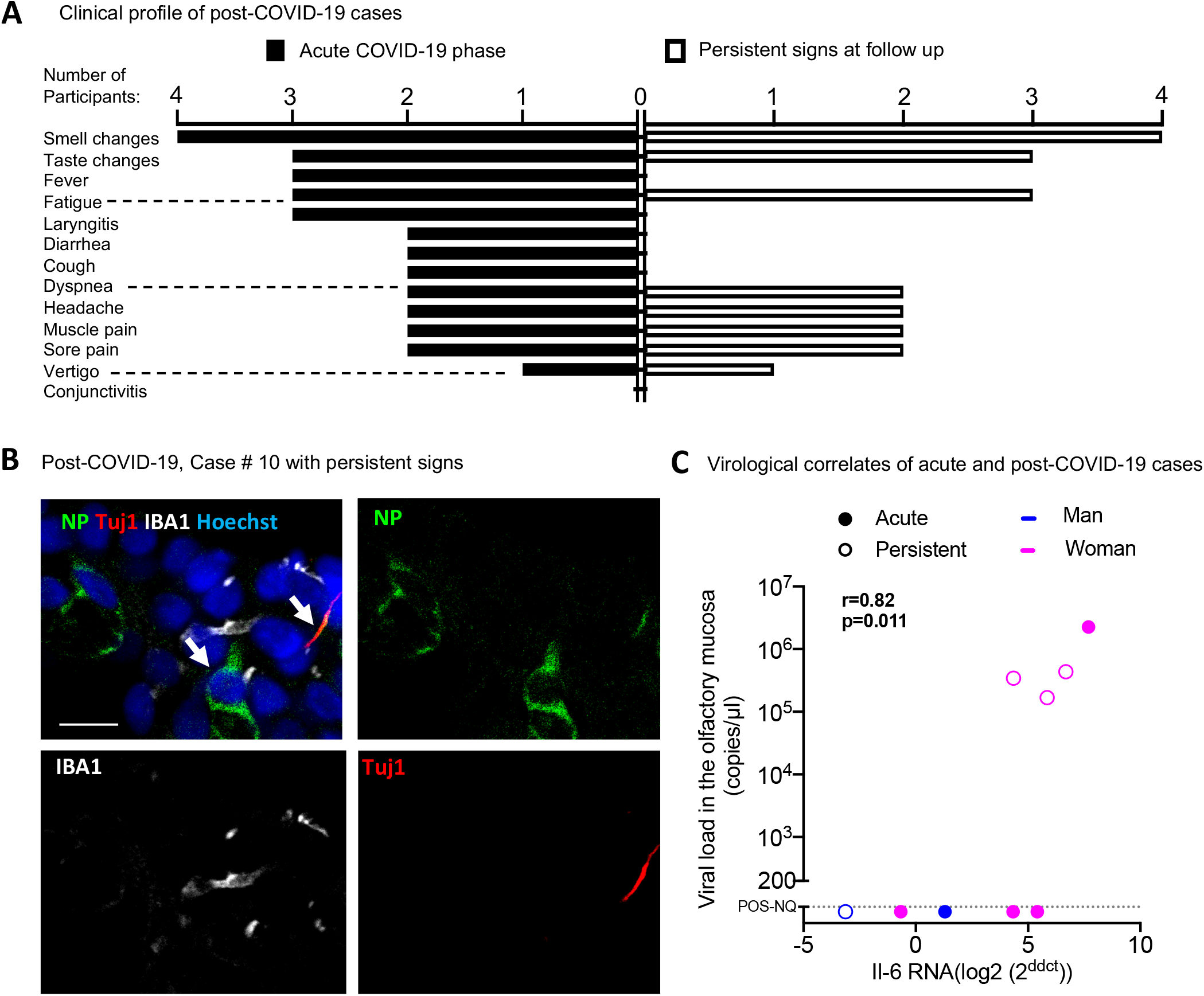
SARS-CoV-2 is present in the olfactory mucosa from patients with persistent loss of smell post-COVID-19. **(A**) Clinical profile of the 4 patients with prolonged loss of smell post-COVID-19. The general symptoms at the acute phase and at the follow up (inclusion in CovidSmell study) are shown. (**B**) Immunofluorescence of infected cells in the olfactory mucosa of the case #10 presenting with persistent olfactory dysfunction at 196 days after COVID-19 onset. The left arrow indicates an infected cell with viral NP staining. The right arrow indicates a Tuj1-NP co-labelling in another cell. (**C**) Graph depicting the correlation between the *IL-6* mRNA expression and the viral load in the 9 patients with acute COVID-19 (“acute”: n=5) or persistent olfactory dysfunction post-COVID-19 (“persistent”, n=4). Viral load values were assessed by Taqman qPCR; when not quantifiable (nq: <200 copies /μL), they were arbitrary given the 50 value for Spearman test (C). Variations in the cytokine gene expression were calculated as the n-fold change in expression in the swabs compared with the swab value of control #1 that was arbitrary put on zero value. Spearman test (C). Scale bar (B): 20 μM.

All had persistent smell loss, persistent taste dysfunction (except case #8) and/or other neurological deficits after COVID-19 at inclusion (Figure 7A) and were seen at the ENT department for this reason. Neurological signs were stereotypical crises of wriggling nose, left intercostal and non-specific arm pain (case #8), paresthesia (case #9) and vertigo (case #10). The characteristics of taste and smell abnormalities at inclusion are described in Supplemental Table S2. Two patients complained of bad taste (Supplemental Table S2). Reduced or increased acuity for bitter, reduced acuity for salt or sour were reported by the two patients with dysgeusia. None of the patients required hospitalization.

No patient had detectable SARS-CoV-2 RNA in nasopharyngeal samples at inclusion during the prolonged phase by the mean of routine diagnosis RT-qPCR. However, all patients had detectable SARS-CoV-2 RNA in olfactory mucosa cytological samples from the olfactory mucosa, using the RT-qPCR SYBR technique (Table 2). Three patients (but not case #6) had a high viral load in the olfactory mucosa (1.68 to 4.35 10^5^ RNA copies/μL; Taqman technique). We further evaluated olfactory mucosa infection by immunofluorescence labeling. We found variable cellularity between olfactory mucosa samples within patients, but all samples contained immature OSNs, positive for Tuj1, indicating the efficient sampling of the neuroepithelium. Immunostaining revealed the presence of SARS-CoV-2 antigens (N protein) in 3 out of 4 patients (Table 2, Figure 7).

We observed abundant Iba1^+^ immune cells in the olfactory mucosa of all patients (Table 2, Figure 7). Quantification of *IL6* gene expression revealed an upregulation of this proinflammatory cytokine in the olfactory mucosa of the 3 patients with high viral load, but not in the case #6, which nevertheless presented SARS-CoV-2 antigens in the neuroepithelium. *IL6* levels in the patients with persistent signs of COVID-19 were similar to those of patients with acute COVID-19 (Tables 1–2, Figure 7C). No changes were observed in *Ccl5, Cxcl10, Isg20* and *Mx1* transcripts. Collectively, these data indicate that the olfactory neuroepithelium from patients with persistent olfactory function loss remains inflamed and infected, with persistent SARS-CoV-2 RNA in all of them.

## Discussion

By combining investigations of COVID19-associated olfactory function loss in patients and experimentally-infected hamsters, both naturally permissive to SARS-COV-2 infection, we demonstrate that multiple cell types of the olfactory neuroepithelium are infected during the acute phase, at the time when loss of smell manifests, and that protracted viral infection in the olfactory neuroepithelium likely accounts for prolonged hyposmia/anosmia.

Strikingly, olfactory mucosa cytological sampling collected from acute or chronically COVID-19 patients with olfactory function loss revealed the presence of the SARS-CoV-2 in 7/9 patients (78%) while the virus was undetected by RT-qPCR performed at inclusion on conventional nasopharyngeal swabs. Therefore, diagnosing SARS-CoV-2 infection in olfactory mucosa sampled by use of nasal cytobrushes might be a more sensitive approach, at least in patients with olfactory function loss, than conventional nasopharyngeal samples. This presence of SARS-CoV-2 RNA and proteins (although the virus infectivity could not be assessed) may influence care management of COVID-19 patients as it may play a role in virus transmission from patients who are thought to be viral-free based on conventional testing, particularly in individuals with mild or no symptoms.

We therefore confirm that SARS-CoV-2 has a significant tropism for the olfactory mucosa (33) and, most importantly, we demonstrate that it can persist locally, not only a few weeks after general symptoms resolution (34–36), but during several months in both mature and immature olfactory sensory neurons. Hence, we found that SARS-CoV-2 persists in the olfactory mucosa of patients with prolonged olfactory function loss, up to 6 months after initial diagnosis. Sampling of the olfactory mucosa revealed viral RNA as well as viral antigens, indicating that long-lasting olfactory function loss in these patients correlates with persistence of both viral infection and inflammation, as shown by high levels of inflammatory cytokines including *IL6,* and the presence of myeloid cells in cytological samples. While reinfection by SARS-CoV-2 could not be formally excluded in these patients (31), the fact that they showed uninterrupted olfactory dysfunction since the onset of the disease, as well as the very low incidence of COVID-19 in France at the time of inclusion, does not support this hypothesis.

To further study anosmia and the inflammatory process in the olfactory system in the context of COVID-19, we used the golden hamster as an animal model for COVID-19. We show that intranasal SARS-CoV-2 inoculation in hamsters leads to infection of OSNs and induces anosmia, accurately recapitulating what is observed in patients, both clinically and histopathologically. Infection of OSNs in SARS-CoV-2-inoculated hamsters has been reported in experiments using similar viral inoculum (26), but not when the inoculum was lower (21), suggesting a dose-dependent susceptibility of OSNs to infection (5, 27, 37, 38). As observed in the tracheal epithelium (39), infection of the neuroepithelium is associated with cilia loss of the OSNs. Once cilia are restored in the late phase of infection *(i.e.,* 14 dpi), olfaction resumes, despite the presence of inflammatory signs. Anosmia thus likely reflects an infection-associated sensorineural dysfunction rather than a simple nostril obstruction or tissue inflammation.

Along with OSNs infection and deciliation, a significant inflammatory process takes place in the nasal cavity and spreads to the olfactory bulb. This inflammatory transcriptional signature, as shown by RNA Seq and confirmed by qPCR for *Il6* but also for *Cxcl10* and *Ifnb1,* is consistent with the recent neuropathological description of deceased COVID-19 patients, where microgliosis was seen in the olfactory bulb (12). Importantly, the fact that similar neuropathological alterations are observed in COVID-19 patients and infected animals implies that SARS-CoV-2 infection is likely the cause rather than a consequence of intensive care provided to COVID-19 patients, as was hypothesized (40).

Although several viruses are known to invade and infect the brain, whether SARS-CoV-2 does so is highly debated. For instance, viral RNA has been detected in the cerebrospinal fluid and other brain tissues collected from patients who died from COVID-19 (12), but the neurological significance of these observations remains unclear (6, 13, 41). The potential SARS-CoV-2 portals of entry to the CNS are (*i*) retrograde neuroinvasion (via olfactory sensory neurons, glossopharyngeal and/or vagal nerve), (*ii*) via the blood-brain barrier endothelium (13, 42) and (*iii*) via peripheral immune cells infiltration (*e.g.*, T-cells and/or peripheral macrophages). Although this does not rule out other routes, our study indicates that SARS-CoV-2 does invade the CNS via the retrograde olfactory pathway. Importantly, in addition to the olfactory bulb, SARS-CoV-2 RNA was also detected in more remote brain areas of infected hamsters, such as the cerebral cortex and the brainstem, yet without clear visualization of viral antigens. Similarly, viral RNA or protein were observed in the brainstem of COVID-19 human patients (12, 37), the location of central cardiorespiratory nuclei. This feature might participate to the pathogenesis of the respiratory distress reported in COVID-19 patients and this study therefore constitutes an important step towards elucidating COVID-19-associated putative neurological dysfunctions. Whether neuronal structures are directly targeted by SARS-CoV-2, as opposed to damage by systemic immune responses, is of particular clinical relevance since these two scenarios would require different therapeutic strategies.

The persistence of long-lasting COVID-19 symptoms is an important topical issue as the pandemic continues (43). Altogether, this work demonstrates a persistent loss of olfactory function in humans with SARS-CoV-2, for multiple months, lasting as long as the virus remains in the same microenvironment. This might result from direct damage to the OSNs which detect odor in the olfactory epithelium. Further, it provides evidence of SARS-CoV-2 retrograde neuroinvasion via the olfactory route leading to neuroinflammation, and shows the association between viral presence in the olfactory epithelium and anosmia, in both acute (hamsters and humans) and long-lasting in COVID-19 patients. The findings we present are clinically relevant in the care to COVID-19 patients, since olfactory function loss could be regarded as a sensitive sign of persistent viral infection, and should be considered in patient management, in particular when antiviral treatments become available.

## Methods

### Patients and study design

Subjects with recent olfactory function loss consulting in the Lariboisière hospital (Paris, France) in the context of the COVID-19 screening care for a suspected or confirmed SARS-CoV-2 infection were included in spring and summer 2020. We also recruited subjects with long-lasting/recurrent loss of smell after COVID-19. Those patients were recruited at the Hotel Dieu Hospital clinic dedicated for long COVID patients. We also recruited in this study control subjects consulting in the Ear, Nose and Throat department at the Lariboisière hospital (Paris, France) with no biologically confirmed COVID-19 or suspected COVID-19 in the past 8 weeks, and no symptoms suggestive of COVID-19 or another respiratory disease and therefore no recent taste and smell function loss. All patients had a detailed standardized clinical and rhinological examination performed by a certified ear nose throat consultant. Following measures were performed at inclusion: *i)* Taste and olfactory function evaluation by a self-questionnaire taste and smell survey (TTS) (44), and a visual analogue scale (VAS) (45), and *ii*) Nasal brushing for collection of neuroepithelium cells and olfactory mucus. The participants self-assessed their smell and taste perception using a 100-mm VAS, where 0 mm indicated the inability to smell or taste and 100 mm indicated normal smell or taste perception (45).

### Human nasal cytobrushes sampling

A certified ear nose throat (ENT) physician sampled olfactory mucosa of each participant by nasal brushing with safety precautions and after local xylocaine application (Lidocaine 5%) following the method previously described (30). Briefly, sampling was performed with a sterile 3.5 mm endocervical brush (02.104, Gyneas, Goussainville, France) inserted and gently rolled five times around the inside of both nostrils (360°). Swabs (one per nostril) were placed on ice immediately following collection, and frozen at −80°C or put in formalin solution 10% neutral buffered (HT-5011-1CS, Sigma).

### Production and titration of SARS-CoV-2 virus

The strain 2019-nCoV/IDF0372/2020 (EVAg collection, Ref-SKU: 014V-03890) was provided by Sylvie Van der Werf, Institut Pasteur, Paris. Viral stocks were produced on Vero-E6 cells infected at a multiplicity of infection of 1.10^-4^ PFU (plaque-forming units). The virus was harvested 3 days post infection, clarified and then aliquoted before storage at −80°C. Viral stocks were titrated on Vero-E6 cells by classical plaque assay using semisolid overlays (Avicel, RC581-NFDR080I, DuPont)(46).

### SARS-CoV-2 model in hamsters

Male and female Syrian hamsters *(Mesocricetus auratus)* of 5-6 weeks of age (average weight 60-80 grams) were purchased from Janvier Laboratories and handled under specific pathogen-free conditions. Hamsters were housed by groups of 4 animals in isolators in a Biosafety level-3 facility, with *ad libitum* access to water and food. Before any manipulation, animals underwent an acclimation period of one week. Animal infection was performed as previously described with few modifications (47). Briefly, the animals were anesthetized with an intraperitoneal injection of 200 mg/kg ketamine (Imalgène 1000, Merial) and 10 mg/kg xylazine (Rompun, Bayer), and 100 μL of physiological solution containing 6.10^4^ PFU (plaque-forming units) of SARS-CoV-2 (strain 2019-nCoV/IDF0372/2020, from Pr Sylvie van der Werf) was administered intranasally to each animal (50 μL/nostril). Mock-infected animals received the physiological solution only. Infected and mock-infected animals were housed in separated isolators and all hamsters were followed-up daily when the body weight and the clinical score were noted. The clinical score was based on a cumulative 0-4 scale: ruffled fur, slow movements, apathy, stress when manipulated. At predefined time-points post-infection, animals were submitted to behavioral tests or euthanized, when samples of nasal turbinates, trachea, lungs and the brain (separated in olfactory bulbs, cerebellum, cortex and brainstem) were collected, immediately frozen at −80°C or formalin-fixed after transcardial perfusion with a physiological solution containing 5.10^3^ U/mL heparin (choay, Sanofi) followed by 4% neutral buffered formaldehyde.

### Behavioral tests

All behavioral assessment was performed in isolators in a Biosafety level-3 facility that we specially equipped for that.

#### Sucrose preference test

We measured taste in hamsters by a sucrose preference test based on a two-bottle choice paradigm which paired 2% sucrose with regular water (48). A reduction in the sucrose preference ratio in experimental infected relative to mock animal is indicative of taste abnormalities. After 6 hours water deprivation, we realized an individual overnight testing which corresponds to a natural activity period of the hamster. The preference was calculated using the following formula: preference = sucrose intake/total intake x 100%. The total intake value is the sum of the sucrose intake value and the regular water intake

#### Buried food finding test

To assess olfaction, we used the buried food finding test as previously described (49) with few modifications. Hamsters were used only once for each test. Four days before testing, Hamsters received chocolate cereals (Coco pops, Kellogg’s) that they ate within one hour. Twenty hours before testing, hamsters were fasted and then individually placed into a fresh cage (37 x 29 x 18 cm) with clean standard bedding for 20 minutes. Hamsters were placed in another similar cage for 2 minutes when about 10-12 pieces of cereals were hidden in 1.5 cm bedding in a corner of the test cage. The tested hamster was then placed in the opposite corner and the latency to find the food (defined as the time to locate cereals and start digging) was recorded using a chronometer. The test was carried out during a 15 min period. As soon as food was uncovered, hamsters were removed from the cage. One minute later, hamsters performed the same test but with visible chocolate cereals, positioned upon the bedding.

### Scanning electron microscopy

For scanning electron microscopy, following animal transcardial perfusion in PBS then 4% neutral buffered formaldehyde, hamster whole heads and lungs where fixed in formalin solution 10% neutral buffered (HT-5011-1CS, Sigma), for one week at 4°C to allow neutralization of the virus. Lung and olfactory epithelium small samples were then finely dissected and post-fixed by incubation in 2.5% glutaraldehyde in 0.1 M cacodylate buffer for 1 h at room temperature then 12 h at 4°C. The samples were washed in 0.1 M cacodylate then several times in water and processed by alternating incubations in 1% osmium tetroxide and 0.1 M thiocarbohydrazide (OTOTO method), as previously described (50). After dehydration by incubation in increasing concentrations of ethanol, samples were critical point dried, mounted on a stub, and analyzed by field emission scanning electron microscopy with a Jeol JSM6700F operating at 3 kV.

### Immunofluorescence

Tissues from PFA-perfused animals were post-fixed one week in PFA 4%, and olfactory brushes from patients were kept in PFA until further use. After post-fixation, hamster whole heads (without skin and lower jaw) were decalcified in TBD-2 (6764003, ThermoFisher) for 3-5 days, then sagitally cut in half and rinsed in PBS. Organs or brushes were then washed in PBS and dehydrated in 30% sucrose. They were then embedded in O.C.T compound (4583, Tissue-Tek), frozen on dry ice and cryostat-sectioned into 20 μm-thick (hamster organs) or 14 μm-thick (brushes) sections. Sections were rinsed in PBS, and epitope retrieval was performed by incubating sections for 20min in citrate buffer pH 6.0 (C-9999, Sigma-Aldrich) at 96°C for 20min, or overnight at 60°C for whole head sections as they are prone to detaching from the slides. Sections were then blocked in PBS supplemented with 10% goat serum, 4% fetal calf serum and 0.4% Triton X-100 for 2h at room temperature, followed by overnight incubation at 4°C with primary antibodies: rat anti-CD11b (1/100, 550282, BD-Biosciences), rabbit anti-SARS-CoV nucleoprotein (1/500, provided by Dr Nicolas Escriou, Institut Pasteur, Paris), mouse anti-OMP (1/250, sc-365818, Santa-Cruz), chicken anti-Iba1 (1/500, 234006, SynapticSystems), mouse anti-Tuj1 (1/250, MA1-118, ThermoFisher). After rinsing, slides were incubated with the appropriate secondary antibodies (1/500: goat anti-rat Alexa Fluor 546, A11081; goat anti-rabbit Alexa Fluor 488, A11034; goat anti-mouse IgG2a Alexa Fluor 546, A21133; goat anti-chicken Alexa Fluor 647, A32933, Invitrogen) for 2 hours at room temperature. All sections were then counterstained with Hoechst (H3570, Invitrogen), rinsed thoroughly in PBS and mounted in Fluoroumont-G (15586276, Invitrogen) before observation with a Zeiss LM 710 inverted confocal microscope.

### RNA isolation and transcriptional analyses by quantitative PCR from Human nasal cytobrushes

Frozen cytobrushes samples were incubated with Trizol (15596026, Invitrogen) during 5 minutes and the total RNA was extracted using the Direct-zol RNAMicroPrep Kit (R2062, Zymo Research). The presence of the SARS-CoV-2 in these samples was evaluated by one-step qRT-PCR in a final volume of 25 μL per reaction in 96-well PCR plates using a thermocycler (7500t Real-time PCR system, Applied Biosystems, Applied Biosystems). Briefly, 5 μL of diluted RNA (1:10) was added to 20μL of Superscript III Platinum One-Step qRT-PCR mix (Invitrogen 11746-100) containing 12.5 μL reaction mix, 0.4 μL 50 mM MgSO4, 1.0 μL superscript RT and 6.1 μL of nuclease-free water containing the nCoV_IP2 primers (nCoV_IP2-12669Fw: 5’-ATGAGCTTAGTCCTGTTG-3’; nCoV_IP2-12759Rv: 5’-CTCCCTTTGTTGTGTTGT-3’) at a final concentration of 1 μM (51). The amplification conditions were as follows: 1 cycle of 55°C for 20 min, 1 cycle of 95°C for 3 min, 50 cycles of 95°C for 15 s and 58°C for 30 s, 1 cycle of 40°C for 30 s; followed by a melt curve, from 60 °C to 95 °C. The viral load quantification in these samples was assessed using a Taqman one-step qRT-PCR, with the same nCoV_IP2 primers and the nCoV_IP2 probe (5’-FAM-AGATGTCTTGTGCTGCCGGTA-3’-TAMRA). Total RNA from human cytobrushes was also reverse transcribed to first strand cDNA using the SuperScript™ IV VILO™ Master Mix (11766050, Invitrogen). To quantify host inflammatory mediators’ transcripts (IL-6, CXCL10, CCL5, Mx1 and ISG20), qPCR was performed in a final volume of 10 μL per reaction in 384-well PCR plates using a thermocycler (QuantStudio 6 Flex, Applied Biosystems). Briefly, 2.5 μL of cDNA (12.5 ng) was added to 7.5 μL of a master mix containing 5 μL of Power SYBR green mix (4367659, Applied Biosystems) and 2.5 μL of nuclease-free water containing predesigned primers (#249900, Qiagen; QuantiTect Primer Assays *IL-6:* QT00083720; *CXCL10:* QT01003065; CCL5: QT00090083;*Mx1:* QT00090895; *ISG20:* QT00225372; and *GAPDH:* QT00079247). The amplification conditions were as follows: 95°C for 10 min, 45 cycles of 95°C for 15 s and 60°C for 1 min; followed by a melt curve, from 60 °C to 95 °C. Variations in the gene expression were calculated as the n-fold change in expression in the tissues compared with the tissues of the control #1.

### RNA isolation and transcriptional analyses by quantitative PCR from Golden hamsters’ tissues

Frozen tissues were homogenized with Trizol (15596026, Invitrogen) in Lysing Matrix D 2 mL tubes (116913100, MP Biomedicals) using the FastPrep-24™ system (MP Biomedicals). Total RNA was then extracted using the Direct-zol RNA MicroPrep Kit (R2062, Zymo Research: olfactory bulb, trachea and nasal turbinates) or MiniPrep Kit (R2052, Zymo Research: lung, brainstem, cerebral cortex and cerebellum) and reverse transcribed to first strand cDNA using the using the SuperScript™ IV VILO™ Master Mix (11766050, Invitrogen). qPCR was performed in a final volume of 10 μL per reaction in 384-well PCR plates using a thermocycler (QuantStudio 6 Flex, Applied Biosystems). Briefly, 2.5 μL of cDNA (12.5 ng) was added to 7.5 μL of a master mix containing 5 μL of Power SYBR green mix (4367659, Applied Biosystems) and 2.5 μL of nuclease-free water with the nCoV_IP2 primers (nCoV_IP2-12669Fw: 5’-ATGAGCTTAGTCCTGTTG-3’; nCoV_IP2-12759Rv: 5’-CTCCCTTTGTTGTGTTGT-3’) at a final concentration of 1 μM. The amplification conditions were as follows: 95°C for 10 min, 45 cycles of 95°C for 15 s and 60°C for 1 min; followed by a melt curve, from 60 °C to 95 °C. Viral load quantification in hamster tissues was assessed by linear regression using a standard curve of eight known quantities of plasmids containing the *RdRp* sequence (ranging from 10^7^ to 10^0^ copies). The threshold of detection was established as 200 viral copies/μg of RNA. The Golden hamsters’ gene targets were selected for quantifying host inflammatory mediators’ transcripts in the tissues using the *Hprt* (hypoxanthine phosphoribosyltransferase) and the *g-actin* genes as reference (Table S3). Variations in the gene expression were calculated as the n-fold change in expression in the tissues from the infected hamsters compared with the tissues of the uninfected ones using the 2^-ΔΔCt^ method (52).

### Transcriptomics analysis in Golden hamsters’ olfactory bulb

RNA preparation was used to construct strand specific single end cDNA libraries according to manufacturers’ instructions (Truseq Stranded mRNA sample prep kit, Illumina). Illumina NextSeq 500 sequencer was used to sequence libraries. The complete RNA-seq analysis approach is described in the Supplemental information.

### Statistics

Statistical analysis was performed using Stata 16 (StataCorp LLC, Texas, USA) and Prism software (GraphPad, version 8, San Diego, USA), with *p* < 0.05 considered significant. Quantitative data were compared across groups using Mann-Whitney non-parametric test. Categorical data was compared between groups using Fisher exact test. Associations between the viral load, the olfactory and taste scores, the cytokine level, and the time from the first disease symptom were estimated with Spearman non-parametric test. In the animal experiences, time to event were analyzed using Kaplan-Meier estimates and compared across groups using the Logrank test. Level of expression of markers at different dpi were compared to the level pre-infection using Kruskal-Wallis followed by the Dunn’s multiple comparison test for unmatched data.

### Study approval

#### Humans

Patients presenting with suspected COVID and recent loss of smell and control subjects were recruited in the CovidSmell study (Study of the Pathogenesis of Olfactory Disorders in COVID-19, ClinicalTrials.gov identifier NCT 04366934). This interventional study received the approval from the ethical committee “Comité de Protection des Personnes SUD EST IV” under reference 20.04.15.64936 and is compliant with French data protection regulations. All the research participants were included at the Lariboisière Hospital, Paris. They received an oral and written information about the research. Informed consent was obtained by the investigator before any intervention related to the research. The Covidsmell study was performed according to the approved protocol.

#### Hamsters

All animal experiments were performed according to the French legislation and in compliance with the European Communities Council Directives (2010/63/UE, French Law 2013–118, February 6, 2013). The Animal Experimentation Ethics Committee (CETEA 89) of the Institut Pasteur approved this study (2020–23) before experiments were initiated.

## Author contributions

GDM, F Lazarini, SL, CH, RH, YM, ER, DS, ML, HB and PML designed research studies; F Lazarini, GDM, SL, F Larrous, VM, VM and LK performed the experiments; F Lazarini, GDM, SL, CH, BV, F Larrous, VM, EK, TC and LK acquired data; F Lazarini, GDM, SL, F Larrous, GG, ML, VM, EK, TC, HB and PML analyzed and discussed data; SW developed behavioral material for hamsters; CH and BV collected olfactory mucosa samples and acquired informed consent; YM supervised statistical analyses; F Lazarini, GDM, SL wrote the manuscript, which was edited by HB, ML and PML. All authors revised and approved the final version of the manuscript.

## Acknowledgments

We thank all participants for volunteering for the clinical study. The human sample from which strain 2019-nCoV/IDF0372/2020 was isolated has been provided by X. Lescure and Y. Yazdanpanah from the Bichat Hospital, Paris, France. We thank Clement Jourdaine, Department of Otorhinolaryngology, Lariboisière Hospital, Paris, France. The clinical research (“CovidSmell”) is sponsored by Institut Pasteur, Paris. We thank Elodie Turc and Laure Lemée, Biomics Platform, C2RT, Institut Pasteur, Paris, France, supported by France Génomique (ANR-10-INBS-09-09), IBISA and the Illumina COVID-19 Projects’ offer. We also thank Anaïs Perilhou, Olivia Chény and Tan-Phuc Bui Van, Clinical Core, CRT, Institut Pasteur, Paris, France. We thank Kurt Sailor, Erwan Poivet, Gabriel Lepousez and Tarek Sharshar for critical reading of the manuscript, Sylvie Van der Werf (National Reference Centre for Respiratory Viruses hosted by Institut Pasteur, Paris) for the SARS-CoV-2 isolate used in this study, and Nicolas Escriou (Innovation lab: Vaccines, Institut Pasteur, Paris) for providing the anti-SARS-CoV-2 nucleoprotein antibody, Marion Berard, Laeticia Breton and Rachid Chennouf for their help in implementing experiments in the Institut Pasteur animal facilities. This work was supported by Institut Pasteur SARS COV2 TASK FORCE (NeuroCovid project), the *Investissements d’Avenir* program managed by the *Agence Nationale de la Recherche* (ANR) under the reference ANR-11-IDEX-0004-02 and ANR-10-LABX-73, the *Agence Nationale de la Recherche* (ANR-15-CE37-0004-01 “SmellBrain”) and the Life Insurance Company “AG2R-La Mondiale”.

Supplemental material: Figures S1-5. Tables S1-3. Additional methods (open field, painted footprints, transcriptomics) and references.

