## Supplemental information for "COVID-19-associated olfactory dysfunction reveals SARS-CoV-2 neuroinvasion and persistence in the olfactory system"


**Supplemental Figures S1-S5:** Pages 2-6

**Supplemental Tables S1-S3:** Pages 7-9

**Supplemental methods:** Page 10

**Supplemental references:** Page 11

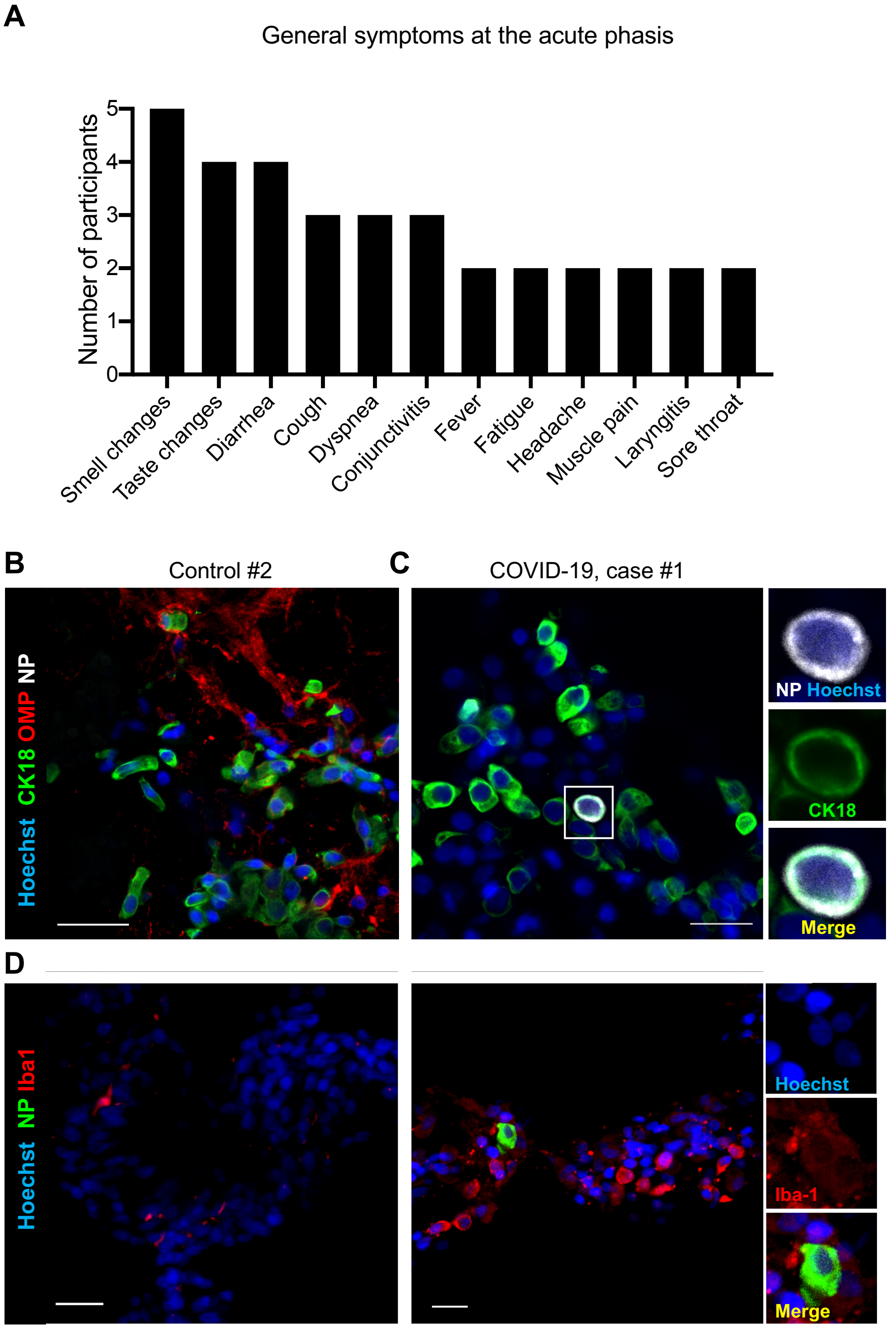


**Figure S1 – General symptoms at the acute phasis and infected cell types in olfactory mucosa of COVID-19 patients at the acute phasis. (A)** Histogram depicting the general symptoms of the COVID-19 patients #1 to #5 at the acute phase. **(B, C)** Immunofluorescence of olfactory mucosa of control (B) and COVID-19 (C) patients, showing olfactory neurons (OMP$^+$ cells) or sustentacular cells (cytokeratin-18 (CK18)$^+$ cells). Inset in (C) shows an infected sustentacular cell. (**D**) Immunofluorescence of COVID-19 patients olfactory mucosa showing infected myeloid cells (Iba1$^+$ cells). Scale bar = 20µm (B, C) or 10µm (D).

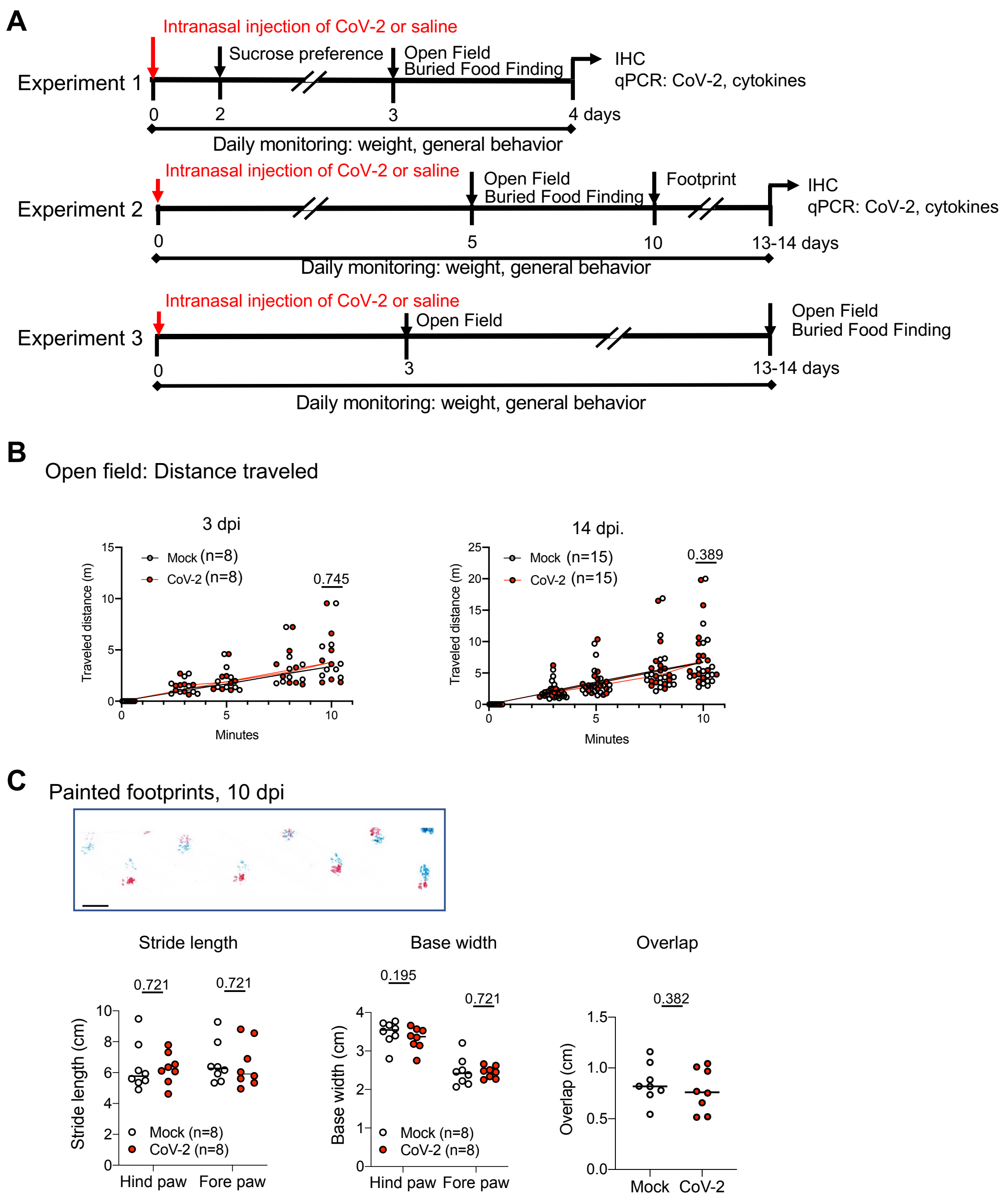


**Figure S2 – Experimental design of the experiments and complementary behavioral tests with golden hamsters.** (**A**) Hamsters were infected intranasally with SARS-CoV-2 or received physiological water (Mock). They were assessed at different timepoints for sensorial, motor and cognitive functions, then terminated at different timepoints for tissue and fluid sampling. Schematic experimental pipeline. (**B**) Total distance traveled in open field at 3 dpi (left) and 14 (right) dpi. (**C**) Analysis of painted footprints leaved by hamsters at 10 days post inoculation, using blue paints on the forepaws and red paints on the hind paws. Up: Picture of footprint patterns. Bottom: Stride length for hind-paw and fore-paw strides (left) Base width hind-paw (middle) and fore-paw steps (right) Overlap between forepaw and hindpaw placement. P values indicated in (A, B, C) are calculated by Mann-Whitney test and are in bold when significant. Scale bar: 5 cm.

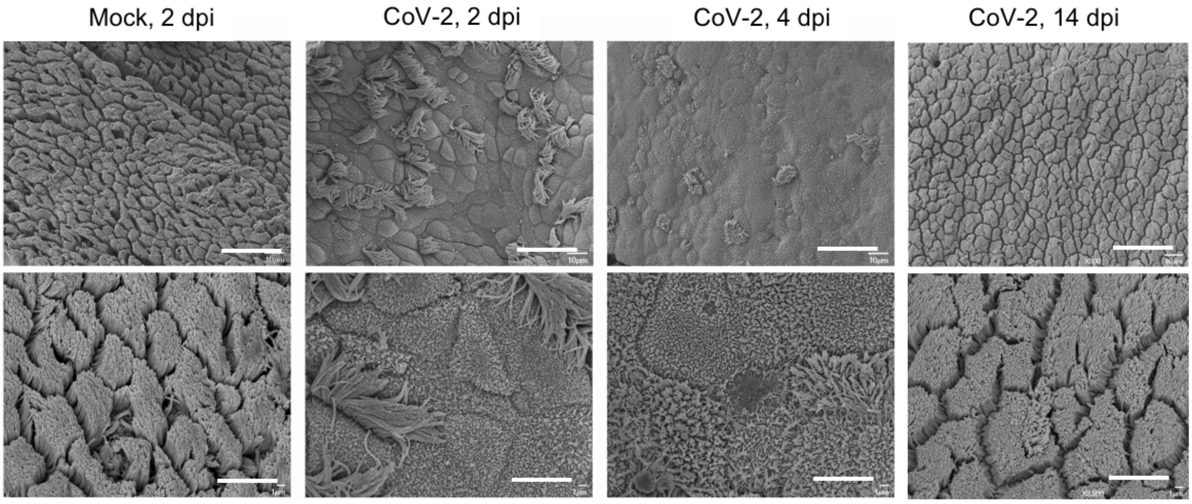


**Figure S3 – SARS-CoV-2 induces loss of ciliation in the olfactory epithelium.** Scanning electron microscope imaging showing changes in olfactory epithelium following CoV-2 infection at 2, 4 and 14 day post intranasal inoculation. Scale bars: 10 µm (up), 1 µm (bottom).

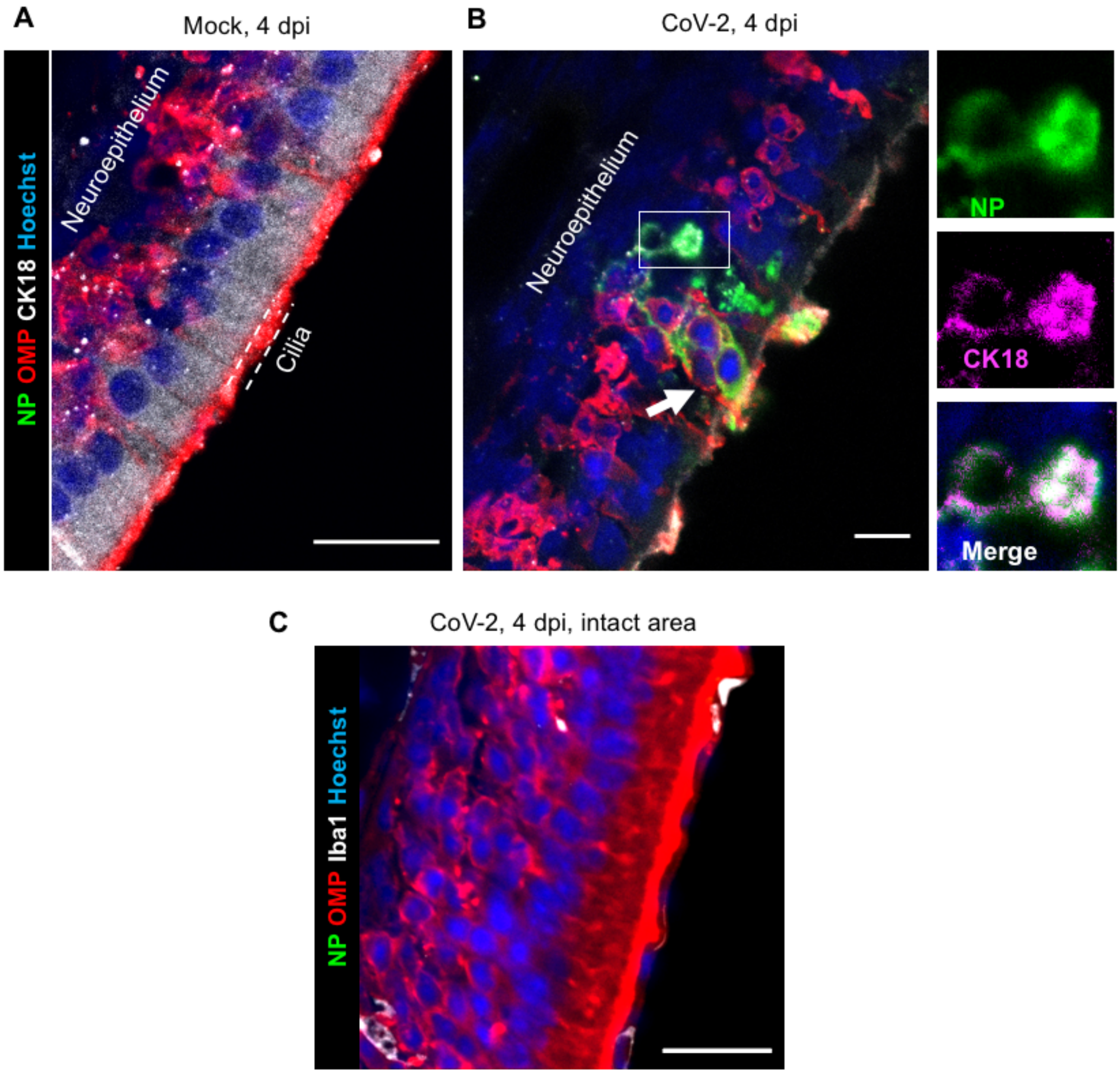


**Figure S4– Cell types infected by SARS-CoV-2 in hamster olfactory mucosa and olfactory bulb.** Olfactory epithelium of mock- (**A**) and SARS-CoV-2 (**B, C**) inoculated hamsters at 4 dpi. Infected sustentacular cells (CK18+) are depicted (inset). Neuro-epithelium containing infected cells are often disorganized (**B**) while adjacent areas without SARS-CoV-2 nucleoprotein staining retained an intact structure (**C**). Mature olfactory neurons express olfactory marker protein (OMP), immature neurons express Tuj1 and myeloid cells express Iba1. SARS-CoV-2 is detected by antibodies raised against the viral nucleoprotein (NP). Scale bars: 20µm.

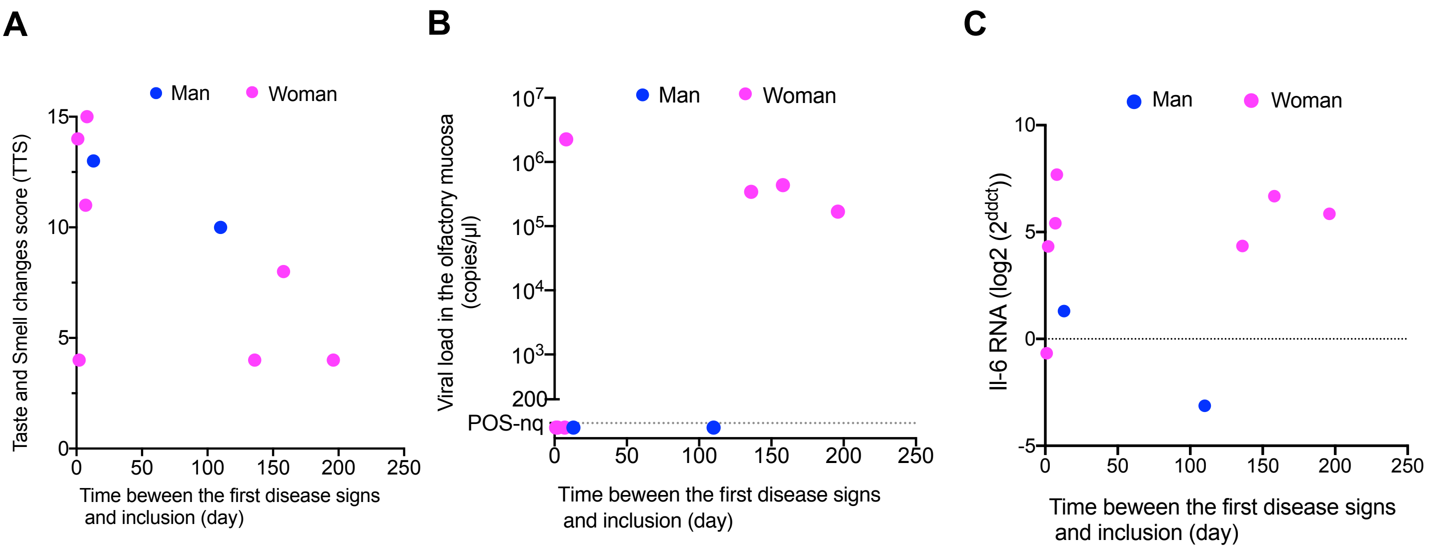


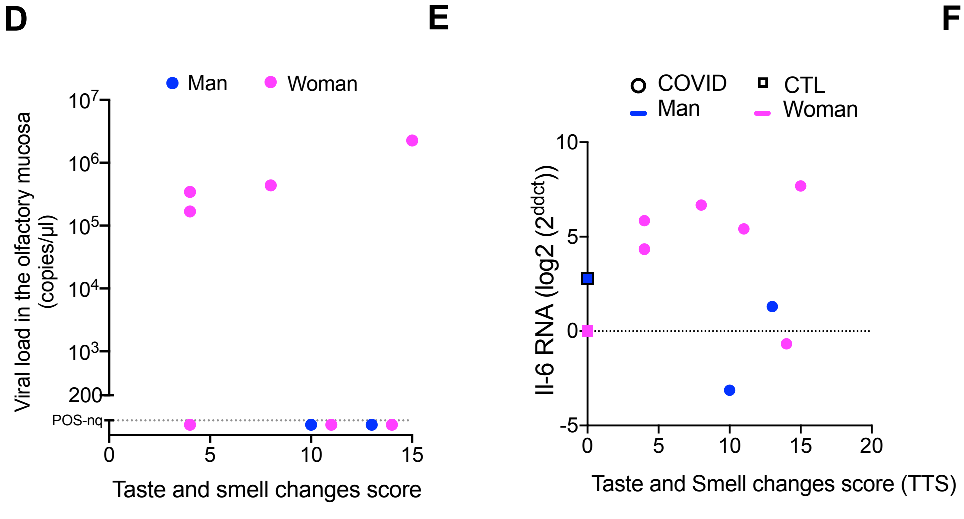


**F**

*r* value

| | |
|---|---|
| 0.80 – 1.00 | Very strong |
| 0.60 – 0.79 | Strong |
| 0.40 – 0.59 | Moderate |
| 0.20 – 0.39 | Weak |
| 0.00 – 0.19 | Very weak |

| R value (p value) | Viral load | IL-6 RNA | TTS |
|---|---|---|---|
| IL-6 RNA | 0.822 (0.011) | | |
| TTS | -0.037 (0.942) | 0.110 (0.747) | |
| Time | 0.530 (0.152) | 0.367 (0.336) | -0.509 (0.169) |

**Figure S5 – Clinical and virologic profiles from patients with persistent olfactory dysfunction post- COVID-19 compared to COVID-19 patients with loss of smell at early onset and controls** Graphs depicting taste and smell changes scores and time between the first disease signs and inclusion (**A**), the viral load in the olfactory mucosa and time between the first disease signs and inclusion (**B**), the level of IL-6 transcripts in the olfactory mucosa and time between the first disease signs and inclusion (**C**), the viral load in the olfactory mucosa and the taste and smell changes scores (**D**), and the taste and smell changes scores and the level of IL-6 transcripts (**E**). TTS: score range 0-15 from 0 no change to 15= anosmia and ageusia.”); n=5 COVID-19 patients with loss of smell included in the early infection phase (“acute”); N=4 patients with persistent olfactory dysfunction post-COVID-19 (“persistent; n=2 control subjects. (**F**) Spearman test r and p are shown.

**Table S1. Characteristics of smell and taste abnormalities at inclusion of the participants with recent loss of smell associated to COVID-19.**

| | Total | COVID #1 | COVID #2 | COVID #3 | COVID #4 | COVID #5 |
|---|---|---|---|---|---|---|
| **Years/Sex** | | 53/W | 31/W | 61/W | 40/W | 46/M |
| **Smell abnormalities** | **5/5** | Yes | Yes | Yes | Yes | Yes |
| Severity of smell loss | | | | | | |
| Partial | 2/5 | Yes | - | Yes | - | - |
| Complete | 3/5 | - | Yes | - | Yes | Yes |
| Reduced acuity | 5/5 | Yes | Yes | Yes | Yes | Yes |
| Increased acuity | 0/5 | No | No | No | No | No |
| « Food smell different » | 2/5 | Yes | Yes | Yes | Yes | Yes |
| Deemed severe | 4/5 | Severe | Severe | Moderate | Severe | Severe |
| First symptom of COVID-19 | 1/5 | No | No | No | No | Yes |
| Preceded of classical symptoms of COVID-19 or minor symptoms | 3/5 | Yes | Yes | Yes | No | No |
| Concomitant with other symptoms of COVID-19 | 1/5 | No | No | No | No | No |
| Sudden onset smell loss | 4/5 | No | Yes | Yes | Yes | Yes |
| Progressive onset smell loss | 1/5 | Yes | No | No | No | No |
| **Taste abnormalities** | **4/5** | Yes | Yes | No | Yes | Yes |
| Severity of Taste loss | | | | | | |
| Partial | 4/4 | Yes | Yes | - | Yes | Yes |
| Complete | 0/4 | - | - | - | - | - |
| Reduced acuity for bitter | 2/4 | No | Partial | - | No* | Partial |
| Reduced acuity for salt | 3/4 | No | Partial | - | Complete | Partial |
| Reduced acuity for sour | 3/4 | No | Complete | - | Complete | Partial |
| Reduced acuity for sweet | 4/4 | Partial | Partial | - | Complete | Complete |
| "Food tastes different " | 4/4 | Yes | Yes | - | Yes | No |
| Deemed severe | 4/4 | Severe | Severe | - | Severe | Severe |
| Bad taste in the mouth | 4/4 | Yes | Yes (bitter) | - | Yes (bitter) | Yes (bitter) |
| Able to discriminate between | | | | | | |
| Two meats | 1/3 | | Yes | - | Yes | No |
| Two vegetables | 1/3 | | Yes | - | Yes | No |
| Two fruits | 1/3 | | Yes | - | Yes | No |
| Meat and fish | 1/2 | *LD* | Yes | - | Yes | No |

*Stronger perception; LD: Lacking data

**Table S2. Characteristics of smell and taste abnormalities at inclusion of the participants with persistent olfactory dysfunction.**

| | Total | COVID #6 | COVID #8 | COVID #9 | COVID #10 |
|---|---|---|---|---|---|
| **Years/Sex** | | 24/M | 43/W | 71/W | 56/W |
| **Smell abnormalities** | **4/4** | Yes | Yes | Yes | Yes |
| Severity of smell loss | | | | | |
| Partial | 0/4 | - | - | - | - |
| Complete | 4/4 | Yes | Yes | Yes | Yes |
| Reduced acuity | 3/4 | Yes | No | Yes | Yes |
| Increased acuity | 0/4 | No | Nos | No | No |
| « Food smell different » | 4/4 | Yes | Yes | Yes | Yes |
| Deemed severe | 1/4 | Severe | Moderate | Moderate | Unimportant |
| First symptom of COVID-19 | 0/4 | No | No | No | No |
| Preceded of classical symptoms of COVID-19 or minor symptoms | 2/4 | LD | Yes | Yes | No |
| Concomitant with other symptoms of COVID-19 | 1/4 | LD | No | No | Yes |
| Sudden onset smell loss | 4/4 | Yes | Yes | Yes | Yes |
| Progressive onset smell loss | 0/4 | No | No | No | No |
| **Taste abnormalities** | **3/4** | Yes | No | Yes | Yes |
| Severity of Taste loss | | | | | |
| Partial | 3/3 | Yes | - | Yes | Yes |
| Complete | 0/3 | - | - | - | - |
| Reduced acuity for bitter | 0/3 | No | -* | No | LD |
| Reduced acuity for salt | 0/3 | No | - | No | No |
| Reduced acuity for sour | 0/3 | No | - | No | No |
| Reduced acuity for sweet | 0/3 | No | - | No | No |
| "Food tastes different " | 3/3 | Yes | - | Yes | Yes |
| Deemed severe | 1/3 | Severe | - | Moderate | Unimportant |
| Bad taste in the mouth | 2/4 | Yes (bitter, sour) | No | Yes | No |
| Able to discriminate between | | | | | |
| Two meats | 0/3 | No | - | No | No |
| Two vegetables | 0/3 | No | - | No | No |
| Two fruits | 0/3 | No | - | No | No |
| Meat and fish | 0/2 | DNK | - | No | No |

DNK: Does not know, LD: Lacking data

**Table S3. Primer sequences used for qPCR in the golden hamster tissues.**

| Gene | | Primer sequence (5' – 3') | Reference |
|---|---|---|---|
| **ha-*γactin*** | For | ACAGAGAGAAGATGACGCAGATAAT | (1) |
| | Rev | GCCTGAATGGCCACGTACA | |
| **ha-*Hprt*** | For | TGCGGATGATATCTCAACTTTAACTG | (2) |
| | Rev | AAAGGAAAGCAAAGTTTGTATTGTCA | |
| **ha-*Il-6*** | For | GGACAATGACTATGTGTTGTTAGAA | (1) |
| | Rev | AGG CAA ATT TCC CAA TTG TAT CCA | |
| **ha-*Cxcl10*** | For | GCCATTCATCCACAGTTGACA | (2) |
| | Rev | CATGGTGCTGACAGTGGAGTCT | |
| **ha-*Ccl5*** | For | ACTGCCTCGTGTTCACATCA | (3) |
| | Rev | CCCACTTCTTCTTTGGGTTG | |
| **ha-*Irf7*** | For | CACTATCCGTGGCTACACTCTG | (3) |
| | Rev | GGTCCTACTCTGTGATGTGCTG | |
| **ha-*Mx2*** | For | CCAGTAATGTGGACATTGCC | (2) |
| | Rev | CATCAACGACCTTGTCTTCAGTA | |
| **ha-*Stat1*** | For | CAATATAGCCGCTTTTCTTTGG | (3) |
| | Rev | TGTACAGGATCCTCCTGGAAGT | |
| **ha-*Ddx58*** | For | CGCGGAACTTTGAAGAGAAG | (3) |
| | Rev | TTGGTCTCCGGCTTTAAGTG | |
| **ha-*Il-1β*** | For | GGCTGATGCTCCCATTCG | (2) |
| | Rev | CACGAGGCATTTCTGTTGTTCA | |
| **ha-*Ifnβ*** | For | ACCCTAAAGGAAGTGCCAG | (4) |
| | Rev | CCAGCTGCCAGTAATAGCTC | |

### Supplemental methods

#### Behavioral tests

Open field: Open Field was employed to check spontaneous locomotor activity. The test was assessed by individually videotracking animals in a 37 x 29 x 18 cm cage with a camera (C920 HD Pro, Logitech) and a single-mouse-tracker (http://icy.bioimageanalysis.org/plugin/single-mouse-tracker/) (5). Total distance performed by the animals during the 10 min of exploration of the arena was recorded.

Painted footprints: In the footprint test, which is used to analyze abnormal gait, the fore- and hind-paws of hamsters were painted in blue and red, respectively, and made to walk straight on a laboratory-made 60 cm length runway (9 x 20 cm) covered with white paper toward a dark goal-box at the end. The footprint patterns obtained on the white paper were then manually analyzed for stride length and base width for both fore- and hind-paws (6).

#### Transcriptomics analysis in Golden hamsters' olfactory bulb

The RNA-seq analysis was performed with the Sequana framework (7). We used the RNA-seq pipeline (v0.9.16), which is available online (https://github.com/sequana/sequana_rnaseq)). It is built on top of Snakemake 5.8.1 (8). Reads were trimmed from adapters using Cutadapt 2.10 (9) then mapped to the golden hamster MesAur1.0 genome assembly from Ensembl using STAR 2.7.3a (10). FeatureCounts 2.0.0 (11) was used to produce the count matrix, assigning reads to features using annotation MesAur1.0.100 with strand-specificity information. Quality control statistics were summarized using MultiQC 1.8 (12). Statistical analysis on the count matrix was performed to identify differentially regulated genes, comparing infected versus non-infected samples considering all samples and separating by sex. Clustering of transcriptomic profiles were assessed using a Principal Component Analysis (PCA). Differential expression testing was conducted using DESeq2 library 1.24.0 (13) scripts based on SARTools 1.7.0 (14) indicating the significance (Benjamini-Hochberg adjusted p-values, false discovery rate FDR < 0.05) and the effect size (fold-change) for each comparison. Finally, enrichment analysis was performed using modules from Sequana, first by converting golden hamster ensembl ids to gene names and then using human annotations for GO terms and KEGG pathways. The GO enrichment module uses PantherDB (15) and QuickGO (16) services; the KEGG pathways enrichment uses gseapy (https://github.com/zqfang/GSEApy/), EnrichR (17), KEGG (18) and BioMart services. All programmatic access to the online web services were performed via BioServices (19).
